# Modulation of biomolecular aggregate morphology and condensate infectivity

**DOI:** 10.1101/2025.07.30.667758

**Authors:** Josephine C. Ferreon, Kyoung-Jae Choi, My Diem Quan, Phoebe S. Tsoi, Cristopher C. Ferreon, Ulas Coskun, Shih-Chu Jeff Liao, Allan Chris M. Ferreon

## Abstract

Neurodegenerative diseases are characterized by pathological aggregates exhibiting distinct morphologies, such as neurofibrillary tangles and dense circular Lewy body-like structures in Alzheimer’s disease, and round hyaline gel-like inclusions and skein-like filaments in amyotrophic lateral sclerosis. However, the mechanisms driving the formation of these diverse morphological structures remain poorly understood. Employing advanced microscopy, including fluorescence lifetime imaging, we investigated condensate aging and aggregation mechanisms of the prion-like domain of hnRNPA1 (A1PrD), a ribonucleoprotein implicated in both disorders. Using a simplified system across various salt and RNA conditions, we demonstrate that homotypic and heterotypic interactions between A1PrD and RNA significantly influence aggregate morphology and amyloid fibril formation, yielding diverse structures including thin fibrils, solid gels, and filamentous starburst aggregates. By tracking aggregate morphogenesis, we observed shifts in fluorescence lifetimes that reflect differences in condensate microenvironments, highlighting distinct homotypic and heterotypic interaction dynamics. Our findings indicate that amyloid fibril formation can initiate within fluid condensates or at the interfaces of solid gels. Moreover, early amyloid-rich fluid starbursts demonstrated the capability to fuse with or recruit younger amyloid-poor droplets, exemplifying prion-like infectivity and accelerating fibril formation. Collectively, our study provides evidence that the interplay between solution composition and the kinetic balance of liquid-liquid phase separation, gelation, and fibrillation contributes to the diverse pathological aggregate morphologies observed in neurodegenerative diseases.

---

The formation of protein aggregates is a hallmark of various neurodegenerative diseases (NDs), characterized by morphological diversity ranging from amorphous aggregates to highly structured fibrillar assemblies. Protein aggregates found in diseased tissues exhibit distinct morphologies. Lewy bodies, prominent in Alzheimer’s Disease (AD), typically manifest as circular structures featuring a dense protein core measuring up to 20-30 µm in diameter(*1*), surrounded by radiating filaments(*2, 3*). Amyotrophic lateral sclerosis (ALS), characterized by progressive neuron degeneration(*4, 5*), presents unique histopathological features such as skein-like inclusions (SLIs) and hyaline inclusions (HI). HI resemble Lewy bodies, appearing as round, glassy structures with central cores ranging from 2 to 20 µm in diameter, often with peripheral halos, while SLIs are composed of thick filamentous bundles extending up to 40 µm in length(*6*). Recently, star-shaped inclusions have been reported for TDP-43 in LATE diseases(*7*), highlighting additional morphological complexity. Despite extensive research, the precise mechanisms underlying these morphological variations remain unclear. Since the seminal study with FUS, it has been proposed that protein aggregation can also occur through liquid-liquid phase separation (LLPS) (*8–10*), wherein a condensed liquid phase separates from the dilute solution(*11–15*). Aging and maturation of these condensates can trigger a liquid-to-solid phase transition. Subsequently, several proteins linked to neurodegeneration, including Tau, HTT, α-synuclein, TDP-43, and hnRNPA1, have been shown to form fibrils under LLPS-promoting conditions(*12, 16–18*). By significantly increasing local protein concentration, LLPS can accelerate the aggregation process(*13, 19, 20*). However, few studies have addressed the relationship between LLPS-mediated aggregation and classical aggregation pathways studied under non-LLPS conditions. Here, we explored hnRNPA1, a ribonucleoprotein implicated in pathological inclusions in AD and ALS, known for its propensity toward LLPS and fibrillar aggregation(*12, 21–25*). We examined how solution conditions affect the formation of hnRNPA1 aggregates, particularly FUS-like starbursts or thin fibrillar morphologies. Using time-lapse confocal 3D microscopy and fluorescence lifetime imaging microscopy (FLIM), we tracked the morphological evolution of hnRNPA1 prion-like domain (A1PrD) condensates. Our observations revealed that liquid gel-like starbursts can ‘infect’ younger condensates through direct coalescence with phase-matched droplets, exacerbating the formation of starburst morphologies. Our findings demonstrate that solution state and composition substantially influence condensate morphology, dynamics, and amyloid formation, emphasizing the critical interplay between condensate fluidity, gelation, and fibrillation in generating diverse aggregate morphologies.

## Solution composition strongly influences the morphology of aggregates formed by hnRNPA1 and its prion-like domain (A1PrD)

Full-length hnRNPA1 contains two RNA recognition motifs (RRMs) and a highly basic, intrinsically disordered C-terminal PrD known to readily undergo aggregation and engage in non-specific, heterotypic interactions with RNA(*12*) (Fig. 1A). Previously, we demonstrated that A1PrD phase separation is sensitive to physicochemical conditions, including pH, protein and NaCl concentrations, and the presence of RNA(*25*). To further elucidate how LLPS correlates with aggregate formation, we systematically examined A1PrD condensation and amyloidogenesis under varying NaCl and RNA, and protein concentrations (Fig. 1B-F, fig. S1). LLPS was monitored via confocal fluorescence microscopy using Alexa Fluor 488-labeled A1PrD (A488-A1PrD, Fig. 1B) and UV light scattering (Fig. 1C).

**Fig. 1.**
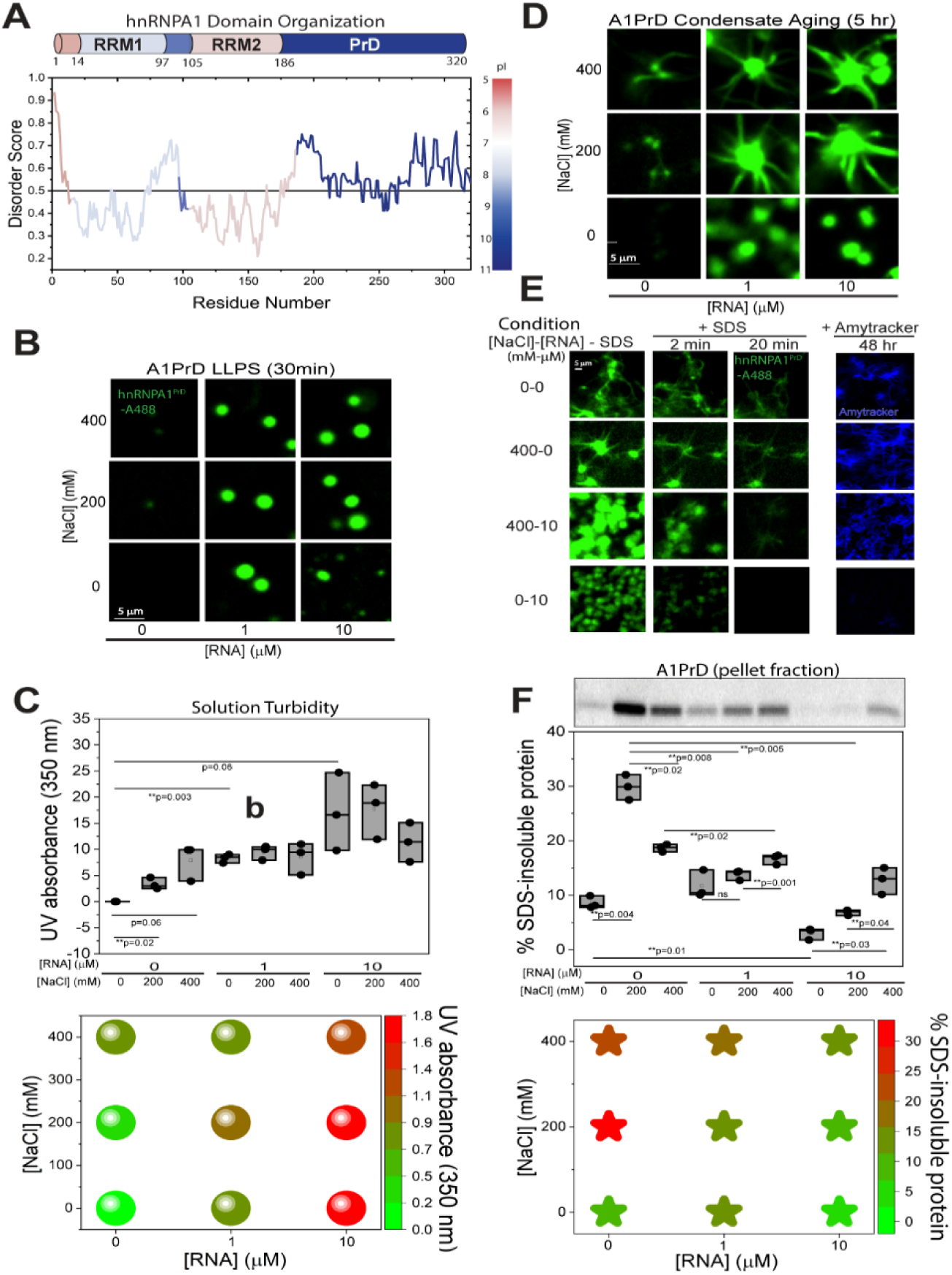
Homotypic and heterotypic interactions modulate A1PrD LLPS and aggregate morphology. **(A)** Domain structure of hnRNPA1 featuring two RNA recognition motifs (RRMs) and a C-terminal prion-like domain (PrD). Disorder prediction was performed using IUPRED2A^1^, with scores above 0.5 indicating increased disorder. The color gradient reflects calculated isoelectric points (pI) of protein segments. **(B)** Confocal fluorescence images of hnRNPA1 PrD (A1PrD; 20 µM unlabeled protein with 100 nM A1PrD-A488) at various concentrations of NaCl and RNA in αβγ buffer (10 mM acetate, 10 mM phosphate, 10 mM glycine, pH 7.5) after a 30-min incubation. Note: Conditions with 0 mM NaCl and 10 µM RNA exhibited significant surface wetting not depicted in images. **(C)** LLPS of A1PrD monitored by UV light scattering at 350 nm (top panel, n=3) at varying concentrations of NaCl and RNA in αβγ buffer. Data are also represented as a NaCl-RNA phase diagram (bottom panel). **(D)** Fluorescence images of samples from (B) following 5-hr incubation. **(E)** Selected conditions ([NaCl]-[RNA] in mM-µM: 0-0, 400-0, 400-10, and 0-10) after 48 hr of aging. Fluorescence images were captured before (first column) and after adding 1% (w/v) SDS (2 and 20 min, second and third columns, respectively; see movies S1-S2 for time-lapse videos of the 0-10 and 400-0 conditions). The final column displays fluorescence images after incubation with Amytracker 480 (blue) in 1% SDS for 90 min. **(F)** Quantification of SDS-insoluble fractions from various [NaCl]-[RNA] conditions after 48 hr, assessed via SDS-PAGE (top panel). Visualization plot illustrates fibril formation intensity (bottom panel; green indicating lowest and red highest intensity). Statistical significance determined by two-sided paired Student’s t-tests; ns=not significant, \**P*<0.05, \*\**P*<0.01.

At the 30-min timepoint, we observed micron-sized (>2 µm) droplets in conditions containing RNA (Fig. 1B). Increasing RNA from 1 to 10 µM enhanced LLPS, consistent with prior studies (*24*). NaCl effects were more complex: in the absence of RNA, increasing NaCl (0 to 400 mM) promoted LLPS, likely due to electrostatic screening of repulsive homotypic interactions between A1PrD molecules. However, at high RNA concentrations (10 µM), elevated salt reduced LLPS, possibly by disrupting favorable heterotypic A1PrD-RNA interactions (Fig. 1C). The condition with 0 mM NaCl and 10 µM RNA (condition 0-10) exhibited the most pronounced LLPS, characterized by droplet fusion and surface wetting. The absence of salt may have preserved strong electrostatic attraction between the positively charged A1PrD and negatively charged RNA. After 5 hr, aggregate morphologies diverged significantly across conditions (Fig. 1D). In most conditions that showed robust LLPS at 30 min, we observed the emergence of “starburst” morphologies (aggregates with a near-spherical dense core and radiating fibrillar extensions), like those previously described for FUS and hnRNPA1/B1 PrDs(*11, 22*). Interestingly, the 0-0 condition (no salt, no RNA) initially showed no detectable condensates at the micron scale, but over 24-48 hr, irregular filamentous aggregates emerged (Fig. 1E), some displaying dense cores reminiscent of starbursts. Unexpectedly, no starbursts were detected in conditions 0-1 and 0-10 (0 mM NaCl with 1 or 10 µM RNA), despite their high LLPS propensity. This suggests that strong heterotypic A1PrD-RNA interactions may inhibit starburst formation. These findings are consistent with models proposing that RNA or other nucleic acids can modulate aggregation by buffering or redirecting the aggregation pathway(*26*), potentially preventing the structured growth associated with starburst morphologies.

To probe the properties of condensate cores and quantify amyloid fibril content, we applied 1% (w/v) SDS, a detergent known to selectively disrupt weak, reversible interactions while sparing more stable, solid-like fibrillar assemblies. Representative conditions from the [NaCl]-[RNA] matrix (0–0, 0–10, 400–0, 400–10) were evaluated. Within 2–20 min of SDS treatment, we observed rapid dissolution of the dense condensate cores or starburst centers, whereas most peripheral fibrils remained intact (Fig. 1E; Movies S1–S2). To independently confirm the presence of amyloid fibrils, we added Amytracker 480 dye at 90 min. Fibril-specific fluorescence revealed substantial amyloid formation in conditions 0-0, 400-0, and 400-10, but not in 0-10, indicating reduced fibrillogenesis under high-RNA/low-salt conditions. Quantitative analysis of SDS-insoluble fractions by SDS-PAGE further supported these findings, with lower levels of detergent-resistant material in the high-LLPS condition (0-10; Fig. 1F and fig. S2). Taken together, these results reveal an anti-correlation between LLPS propensity and amyloid fibril formation (compare Fig. 1C and 1F, bottom panels; fig. S2), suggesting that highly dynamic liquid-like condensates may buffer or delay the transition to irreversible fibrillar structures.

To characterize microenvironmental differences within condensates, we employed fluorescence lifetime imaging microscopy (FLIM), a robust technique for detecting dynamic changes in local environments associated with macromolecular condensation (*27–29*) and aggregation (*30*) (fig. S3–S5). Longer fluorescence lifetimes typically reflect greater molecular mobility and a more fluid environment, while shorter lifetimes suggest quenching due to increased molecular interactions, crowding, or the formation of dense structures (*27, 28, 30–33*). Under dilute, non-LLPS conditions (∼100 nM A1PrD-A488 without excess unlabeled protein), we observed marked decreases in fluorescence lifetimes upon RNA addition. Lifetimes decreased from ∼3.5 ns in the absence of RNA (conditions 0-0 and 400-0) to ∼3.2 ns at 400-10, and most dramatically to ∼1.5 ns in the 0-10 condition (no salt, high RNA), indicating strong heterotypic A1PrD-RNA interactions (Fig. 2A, top row). These effects were partially mitigated by the presence of salt, which likely screens electrostatic interactions. LLPS was induced by adding 20 µM unlabeled A1PrD, resulting in droplet formation under all conditions except 0-0. FLIM images were collected at early (<2 hr) and late (>5–24 hr) timepoints to monitor the evolution from liquid droplets to starbursts and aggregates. Condensed-phase droplets formed within 2 hr exhibited distinct lifetime shifts relative to dilute conditions. Most notably, lifetimes further decreased to ∼2.2–2.8 ns for all conditions except 0-10, where lifetime increased to ∼2.7 ns, possibly due to buffering of RNA interactions by excess protein (Fig. 2A, middle row). These trends were consistent with partitioning data: higher LLPS propensity (Fig. 2A) correlated with greater protein and RNA enrichment within condensates (Fig. 2B and 2C). In the 0-0 condition (<2 hr), the moderate lifetime reduction may reflect the presence of nano-oligomers or early clustering not visible by microscopy(*24, 34*). At later stages (>5–24 hr; Fig. 2A, bottom row), fluorescence lifetimes remained largely unchanged except in the 0-0 condition, which showed significantly reduced lifetimes (∼1.5 ns) and the appearance of fibrillar aggregates. TEM analysis corroborated these findings, revealing fine, thin filaments in the 0-0 condition and thicker, bundled fibrils in high-LLPS conditions (Fig. 2D). Additionally, we observed ∼100 nm spherical nanocondensates coating many of the fibrils. Interestingly, the 0-10 condition, which lacked starburst formation, contained condensates with small, globular gel-like inclusions (see Fig. 2A, orange frame). We hypothesize that significant gelation occurs during the aging of condensed droplets. This was validated by fluorescence recovery after photobleaching (FRAP) experiments across timepoints: early-stage droplets showed rapid recovery, while aged gels and starbursts exhibited slowed or absent recovery, indicative of a transition to an arrested or solid-like state (Fig. 2E, fig. S7). Altogether, our fluorescence lifetime imaging microscopy (FLIM) data clearly highlights distinct solution microenvironments among different condensate populations. Specifically, these differences align consistently with pathological condensates characterized by homotypic interactions and physiological condensates involving heterotypic interactions with RNA. This distinction underscores the critical role of molecular composition and interaction dynamics in determining the functional and pathological outcomes of biomolecular condensates.

**Fig. 2.**
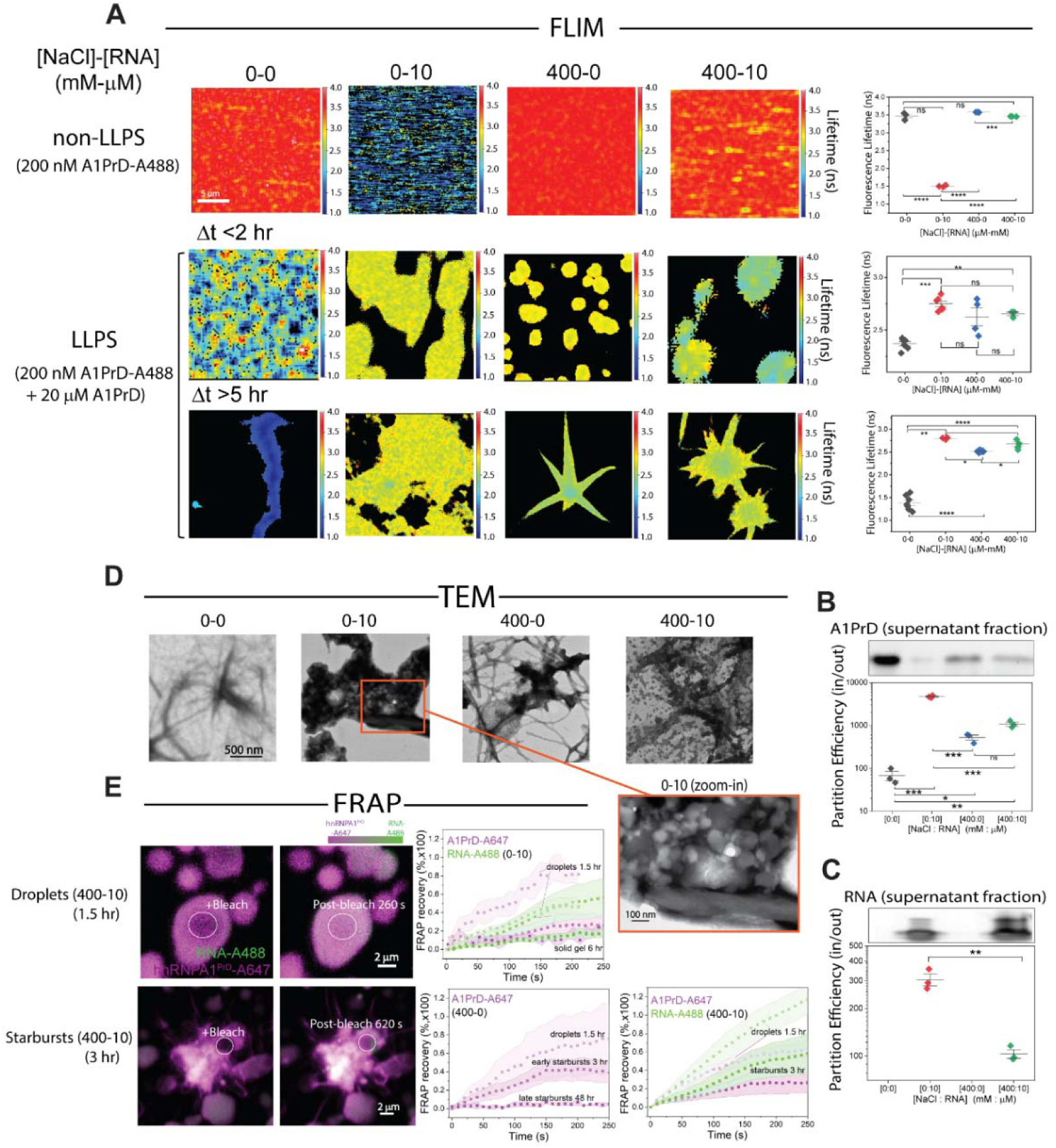
Distinct microenvironments within homotypic and heterotypic condensates. **(A)** Fluorescence lifetime imaging microscopy (FLIM) of A1PrD under selected NaCl-RNA conditions ([NaCl]-[RNA], mM-µM: 0-0, 0-10, 400-0, and 400-10), comparing non-LLPS controls (without unlabeled protein) with LLPS samples at early and late timepoints. Corresponding intensity images are provided in fig. S6. Right panels show quantified FLIM lifetimes (mean ± SD; n=3–7 images per condition). **(B)** Partition efficiency of A1PrD in condensates (inside vs. outside) quantified by SDS-PAGE analysis of supernatant fractions across varying NaCl-RNA conditions (top panel; n=3). **(C)** RNA partition efficiency into condensates, assessed by native gel electrophoresis quantification of supernatant fractions (top panel; n=3). **(D)** Transmission electron microscopy (TEM) images of selected samples from (A) incubated for 48 hr. The zoomed inset (orange frame) for the 0-10 condition highlights gel-like globules within condensates and bundled fibrils at the condensate edges. **(E)** Representative FRAP (Fluorescence Recovery After Photobleaching) images for droplets and starbursts (A1PrD-A647 and RNA-A488; condition 400-10). Right panels show FRAP recovery plots with average fluorescence intensity (symbols) and standard deviation (shaded areas) for A1PrD-A647 (purple) and RNA-A488 (green) across different conditions (400-0, 0-10, and 400-10) and aging times. Representative FRAP images are also shown in fig. S7. Sample sizes: 400-0: droplets n=7, early starbursts n=3, late starbursts n=3; 400-10: droplets n=9, starbursts n=3; 0-10: droplets n=4, solid gels n=5. Darker symbols and shaded areas represent longer-aged samples. Statistical significance determined by two-sided paired Student’s t-tests; ns=not significant, \**P*<0.05, \*\**P*<0.01, \*\*\**P*<0.005, \*\*\*\**P*<0.001.

## A1PrD droplet-to-starburst morphogenesis in the presence of RNA

To elucidate the mechanism of starburst formation, we employed 4D confocal fluorescence microscopy (time-lapse 3D z-stacks) to monitor the transformation of A1PrD liquid droplets into starburst aggregates. We focused on conditions known to robustly promote starburst formation—specifically, 10 µM RNA and 200 mM NaCl. To visualize amyloid formation, Thioflavin T (ThT), a fluorescent dye specific for β-sheet-rich fibrils, was added to the imaging solution. Immediately following LLPS initiation, A1PrD-A647 droplets formed rapidly, exhibiting hallmark liquid behaviors such as fusion and surface wetting within the first hour (Fig. 3A, fig. S8). Within a few hours, we observed the emergence of ThT-positive condensates (orange signal; Fig. 3A and fig. S8, white arrows), which we interpret as amyloid seed nuclei. These seeds progressively enlarged and developed filamentous protrusions over time (Fig. 3A, fig. S8). By 12 hr, the majority of condensates had matured into starbursts (Fig. 3A, Movie S3). Early-stage protrusions (<3 hr) ranged from 0.5–1.2 µm in length, while later-stage structures (>5 hr) extended to ∼5-17 µm. Transmission electron microscopy (TEM) of early droplets revealed a network of fine fibrillar structures (Fig. 3B). Consistent with our earlier observations (Fig. 2E; 400-10 RNA condition), TEM of mature starbursts revealed a chaotic meshwork of thick, coarse filaments (∼200 nm diameter) decorated with nanometer-sized beads or nanocondensates (Fig. 3B). These adsorbed nanocondensates may represent early-stage fibrillar precursors or serve as secondary nucleation sites that facilitate fibril elongation or fragmentation, contributing to the intricate and dynamic architecture of starburst morphologies. Consistent with Fig. 2A, FLIM analysis revealed no dramatic shift in average fluorescence lifetimes between liquid droplets and starbursts, suggesting that the overall microenvironment within condensates remains relatively stable during morphological evolution (Fig. 3C). However, we detected a broadening in the lifetime distribution and a modest shift in the central peak from ∼2.6 to ∼2.9 ns, which was counterintuitive (Fig. 3D). To enable quantitative analysis across multiple datasets, we binned lifetime values into four categories: <2.1 ns, 2.1-2.45 ns, 2.45-2.8 ns, and >2.8 ns (Fig. 3D, right panel). Corresponding phasor plots were generated to map spatial localization of lifetime clusters (figs. S9–S10). Intriguingly, higher lifetime populations predominantly localized to the starburst periphery, which we hypothesize represent newly adsorbed nanocondensates from the dilute phase, structures also observed in TEM analysis (Fig. 3B).

**Fig. 3.**
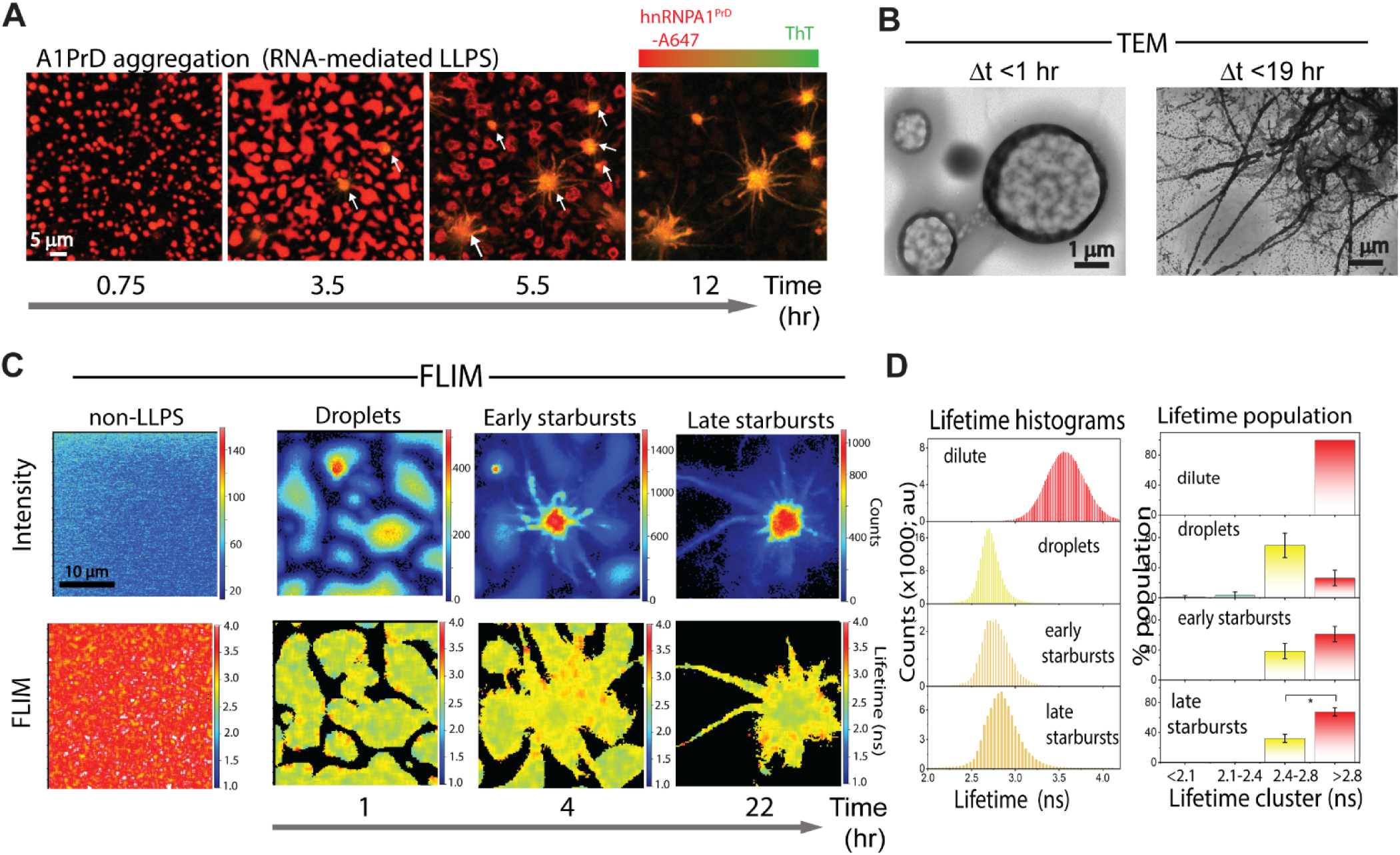
RNA-facilitated aging of A1PrD droplets yields dynamic, fluid starbursts. **(A)** Time-dependent initiation and maturation of A1PrD starbursts monitored by confocal fluorescence microscopy (20 µM A1PrD, 10 µM RNA, ∼70 nM A1PrD-A647 (red), and 3 µM thioflavin T (ThT, green) in αβγ buffer containing 200 mM NaCl). **(B)** TEM images depicting an early-stage condensate (1 hr) and a mature starburst (19 hr). **(C)** Representative confocal (top) and FLIM (bottom) images showing different stages of A1PrD droplet maturation and starburst formation. Conditions include non-LLPS dilute controls and LLPS conditions at 1, 4, and 22 hr post-initiation. **(D)** Fluorescence lifetime histograms corresponding to the images shown in (C). Images were categorized by incubation time and morphological state: surface-wetted droplets (∼1 hr, n=4 images), early starbursts (<5 hr, n=7 images), and late starbursts (>12 hr, n=5 images). Right panels present bar graphs showing lifetime distributions clustered into four bins: <2.1 ns (blue), 2.1–2.45 ns (blue-green), 2.45–2.8 ns (yellow), and >2.8 ns (red). Error bars represent standard deviation (SD) across multiple images (*n indicated above*).

To further investigate starburst growth dynamics, we tracked individual structures over time and performed morphometric quantification using surface rendering in Imaris. We observed that initial starburst nuclei often emerged from above the wetted surface (Fig. 4A, white arrow). As starbursts expanded and filaments elongated, a concurrent depletion of surface-bound liquid droplets was noted (Fig. 4A–B, movies S4-6). Quantitative correlation analysis revealed that increasing ThT fluorescence intensity closely paralleled starburst maturation, including increased surface roughness and filament extension (Fig. 4C-E; movies S4-S6; figs. S11-13). Importantly, early-stage starbursts retained fluid properties, as evidenced by their ability to fuse with other droplets or starbursts. We also observed condensation or shrinkage of some droplets, likely reflecting gelation or adsorption into starburst filaments. These findings align with FLIM data showing heterogeneous lifetime distributions across starburst structures, supporting a model in which each starburst is a multi-nucleated assembly formed through successive fusion and maturation events (Fig. 3C; figs. S9-10).

**Fig. 4.**
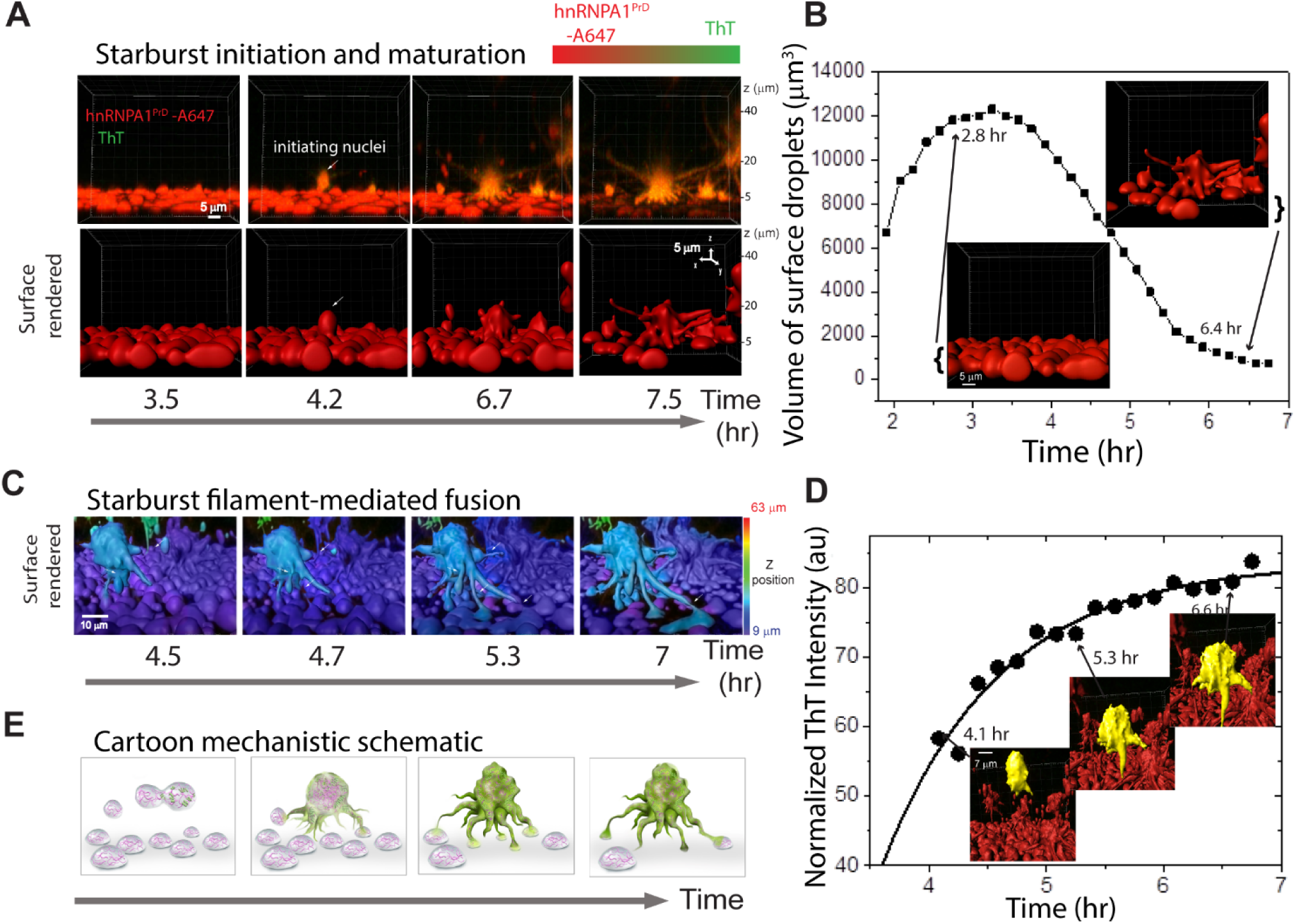
A1PrD starbursts siphon from and infect young condensates. **(A)** Time-dependent maturation of starburst aggregates of 20 µM A1PrD in the presence of 10 µM RNA and 200 mM NaCl (with ∼70 nM A1PrD-A647 and 3 µM ThT in αβγ buffer), visualized by 4D confocal microscopy (time-lapse 3D z-stack). Additional images provided in fig. S8 and movies S4-S5. Bottom panels: Surface-rendered starburst images at various maturation stages generated using Imaris software (movie S6). **(B)** Quantitative plot of the volume of surface-rendered droplets over time. Representative surface-rendered images at selected time points are displayed. **(C)** Surface-rendered visualization depicting fusion events and subsequent in-phase siphoning between starburst filaments and nearby droplets (movie S7). The primary starburst is highlighted in cyan for clarity. **(D)** Temporal plot of normalized ThT fluorescence intensity from tracked starburst aggregates. Representative surface-rendered images (yellow) at specific time points are shown. Similar analyses were performed for five maturing starbursts (fig. S7). **(E)** Schematic illustration of the infection mechanism, wherein ThT-positive (amyloid-rich, green) starburst aggregate siphons material from and recruit ThT-negative condensates.

Upon closer analysis, we observed that the loss of surface-bound droplets was accelerated by a process resembling ‘in-phase siphoning,’ whereby dynamic filamentous extensions of starbursts actively recruited nearby droplets into the growing structure (Fig. 4B; movie S6). Notably, ThT-positive, amyloid-rich starbursts were capable of recruiting ThT-negative liquid droplets. This observation led us to hypothesize that starbursts exhibit a form of ‘prion-like’ seeding behavior, wherein ThT-positive condensates can convert or “infect” ThT-negative droplets upon contact (Fig. 4E; fig. S12; movie S7).

Interestingly, we observed similar starburst morphogenesis in the absence of RNA (movie S8), although fewer LLPS droplets were formed under these conditions. As a result, fusion events between droplets and starbursts were reduced, and the resulting starbursts had smaller core diameters compared to RNA-rich conditions (compare 400-0 and 400-10 in Fig. 2A). These data suggest that filamentation primarily propagates outward from the interior of starbursts. However, we also documented cases where filamentous protrusions appeared to originate from the condensate interface, particularly at the boundary of arrested or gel-like condensates (fig. S14, movie S8), consistent with prior descriptions of interface-driven surface nucleation(*18, 22*). We propose that the capacity of starbursts to undergo both filament elongation and droplet fusion depends on their retention of a fluid and dynamic internal environment during early stages. This is supported by FRAP experiments, which demonstrated rapid fluorescence recovery in early-stage starbursts (Fig. 2E; fig. S7), confirming their liquid-like character and compatibility with dynamic remodeling and recruitment processes.

## Starburst evolution of A1PrD in the presence of a crowding agent

Crowding agents such as polyethylene glycol (PEG) are widely used to promote LLPS (*11, 12*). To compare the mechanisms underlying NaCl-/RNA-facilitated versus crowding agent-mediated starburst formation, we performed time-lapse confocal microscopy tracking under conditions containing 10% (v/v) PEG-8K. PEG-induced LLPS yielded sparser droplets, enabling the tracking of individual starburst evolution events (Fig. 5A, fig. S15, movie S10). The morphogenesis from droplet to starburst closely resembled that previously described for FUS(*11*), characterized by thickening and elongation of radial filaments. TEM analysis of early condensates revealed the presence of oligomeric intermediates and nascent fibrils that initiated from within the condensates and extended outward (Fig. 5B). FLIM analysis revealed distinct differences in fluorescence lifetime patterns compared to RNA-mediated condensates (Figs. 5C–E). As expected, dilute A1PrD solutions (∼70 nM A1PrD-A488) exhibited high lifetimes (∼3.5 ns; Figs. 5C–E; phasor plots in figs. S16-18). Applying the same binning strategy for lifetime clusters (<2.1, 2.1–2.45, 2.45–2.8, >2.8 ns), early PEG-induced droplets (∼30 min) displayed at least two dominant lifetime populations (∼2.2 and ∼2.5 ns; Fig. 5D–E). Notably, fusion events were observed between droplets with distinct lifetime signatures (Fig. 5C, early droplets panels), indicating heterogeneity in their internal environments. With continued incubation, droplets matured into structures exhibiting lifetimes <2 ns (Figs. 5C-E), with some transitioning into starbursts (Fig. 5C, fig. S15). As seen in RNA-induced starbursts, the filamentous extensions and peripheral shell of PEG-induced starbursts retained longer lifetimes (∼2.2-2.5 ns), whereas their dense core exhibited shorter lifetimes (<2 ns), suggestive of increased local quenching due to denser molecular packing (Figs. 5C-E, figs. S16-18). A gradient in lifetimes from the outer interface to the center core was consistently observed, reflecting spatial variation in local microenvironments. Given the high unlabeled-to-labeled protein ratio (∼300:1) and uniform fluorescence intensity across droplets, the observed lifetime reductions likely indicate increased fluorophore quenching driven by molecular crowding in the core. The gradual time-dependent decrease in lifetimes and the emergence of spatial gradients are consistent with prior studies suggesting that condensates behave as aging Maxwell fluids(*35*), transitioning gradually from liquid-like to gel-like or solid-like states rather than undergoing abrupt phase changes. We speculate that the longer lifetimes at droplet and starburst interfaces result from dynamic exchange with the surrounding dilute phase, which averages local environments over time. Furthermore, these interfaces may function as scaffolds or ‘magnets’ for ongoing nucleation and fibril elongation, as similarly observed in RNA-induced starbursts. Altogether, PEG-mediated starburst formation appears to involve three key features: (1) fibrillation originating internally and propagating outward, (2) droplet aging via gelation or glass-like dynamics, and (3) secondary fibril nucleation and branching at interfaces and filament tips.

**Fig. 5.**
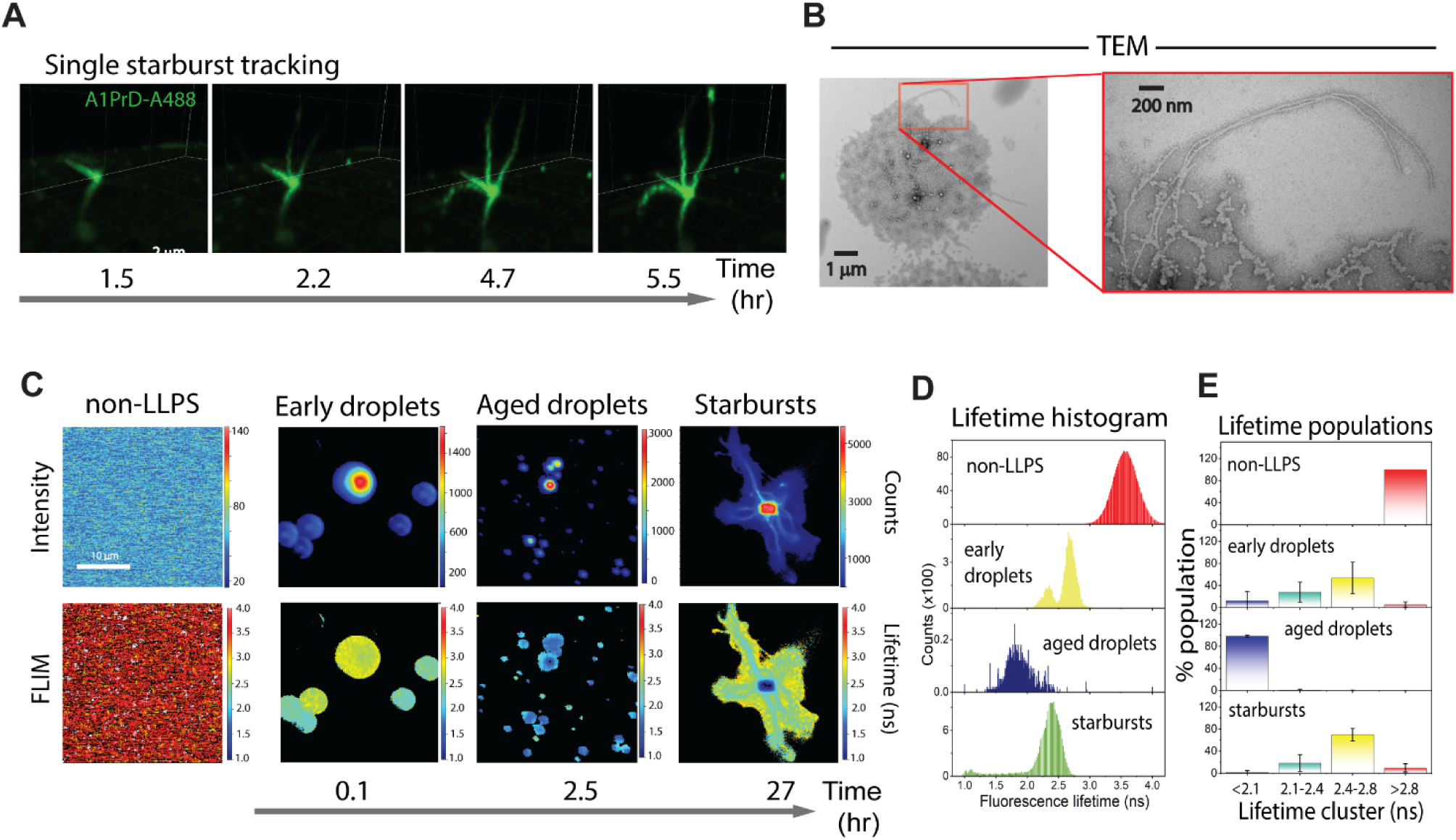
A1PrD droplet aging and starburst formation in the presence of crowding agent (PEG). PEG-enhanced protein condensation was monitored using 20 µM unlabeled A1PrD, ∼70 nM A1PrD-A488, 10% w/v PEG-8K, and 200 mM NaCl in αβγ buffer. **(A)** Time-dependent initiation, maturation, and fibrillar extension of A1PrD starbursts visualized by 4D confocal microscopy (time-lapse 3D z-stack). **(B)** TEM image of an aged condensate showing fibrillar extensions. **(C)** Representative confocal (top row) and FLIM (bottom row) images at different stages of droplet maturation and starburst formation: non-LLPS dilute controls, and LLPS conditions at ∼5 min, 2.5 hr, and 27 hr post-LLPS initiation. **(D)** Fluorescence lifetime histograms corresponding to the images shown in (C). **(E)** Lifetime data grouped according to incubation time and morphological states: early droplets (10–30 min, n=3 images, >200 droplets); aged droplets (2–3 hr, n=4 images, >200 droplets); and mature starbursts (n=7 images). Lifetimes were clustered into four bins: <2.1 ns (blue), 2.1–2.45 ns (blue-green), 2.45–2.8 ns (yellow), and >2.8 ns (red). Error bars represent the average and SD from multiple images (n indicated).

## LLPS vs. non-LLPS aggregation mechanisms

Traditional protein aggregation studies are typically conducted under dilute conditions and proceed via mechanisms involving protein misfolding or the nucleation of fibrillation-competent species (*36*). To investigate A1PrD aggregation under non-LLPS conditions, we employed temperature modulation to reverse LLPS and bypass phase separation-mediated pathways (*37, 38*). A1PrD condensates formed via RNA-mediated LLPS are reversible upon temperature elevation or treatment with the chemical disruptor 1,6-hexanediol (*12, 18, 39*). In contrast, aged starbursts remain resistant to such perturbations (Fig. 6A, movie S11, fig. S19). We utilized this property to induce aggregation via a non-LLPS route by raising the temperature to 55°C, which led to rapid droplet dissolution and allowed us to observe A1PrD aggregation from the dilute phase. Within 30 minutes, we detected small, irregularly shaped aggregates by fluorescence microscopy and TEM (Fig. 6B-C). These microscopic aggregates likely originate from oligomeric intermediates or cluster-like nuclei, which gradually grow into larger irregular structures over several hours. However, even after prolonged incubation, non-LLPS aggregates remained markedly smaller than those formed via LLPS pathways (e.g., PEG- and RNA-mediated conditions; Fig. 6D). TEM analysis revealed that non-LLPS aggregates were composed of fine, thin filaments (Fig. 6C), resembling those observed under low-salt and RNA-free conditions (0-0; Fig. 2E). In contrast, LLPS-mediated aggregates were composed of thicker, coarser filaments (Fig. 6C), underscoring a mechanistic divergence between aggregation pathways. These findings highlight the distinct morphologies and assembly dynamics that result from LLPS-mediated versus non-LLPS aggregation processes.

**Fig. 6.**
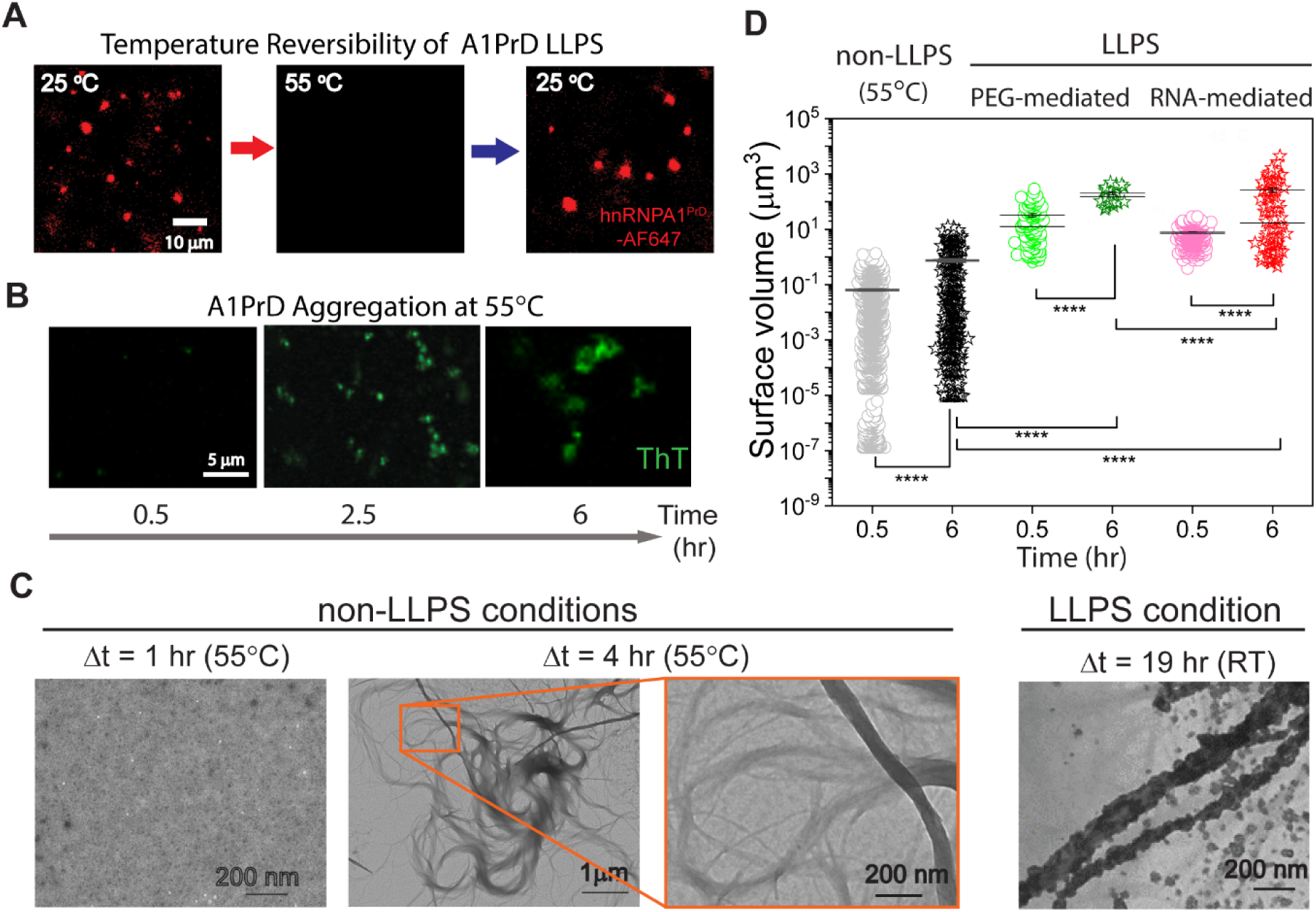
Morphological distinctions between aggregates formed under non-LLPS and LLPS conditions. **(A)** Fluorescence microscopy data demonstrating temperature-dependent reversibility of RNA-mediated A1PrD LLPS droplets (10 µM RNA, 20 µM A1PrD, ∼70 nM A1PrD-A647, and 3 µM ThT in αβγ buffer). **(B)** Confocal fluorescence images showing A1PrD aggregates formed at elevated temperature (55°C) under identical solution conditions as described in (A). **(C)** TEM images illustrating A1PrD particulate and fibrillar aggregates obtained by incubating samples (10 µM RNA, and 20 µM A1PrD in αβγ buffer) at 55°C for 1 hr (particles) and 4 hr (fibrils). **(D)** Comparative size distributions of particle aggregates formed under non-LLPS versus LLPS conditions, facilitated by either PEG or RNA, at incubation times of 30 min or 6 hr.

## Interplay between LLPS, gelation, and fibrillation governs condensate morphogenesis

To unify our findings and provide a conceptual framework for how different physicochemical processes shape condensate morphology, we developed a schematic model depicted in Fig. 7. This model illustrates how the relative kinetics of LLPS, gelation, and fibrillation govern the resulting aggregate structures of A1PrD and similar prion-like proteins. If the kinetics of gelation outpace fibrillation, arrested or gel-like condensates emerge, as observed for A1PrD in the presence of RNA. These arrested states lack pronounced filamentous extensions and often show limited internal restructuring. In contrast, when LLPS and fibrillation proceed more rapidly than gelation, large starbursts form with prominent filamentous protrusions and expanded core diameters, indicative of continued internal fluidity that facilitates outward fibril growth. When gelation is faster than fibrillation but occurs after initial LLPS, fibrils may nucleate and grow from the droplet interface, yielding more solid-like droplets with small dense cores and interface-localized filamentation. Alternatively, if gelation interferes with droplet fusion during LLPS, smaller core structures may form, often manifesting as bead-like globules. Finally, under non-LLPS conditions, such as those induced by elevated temperature or the absence of salt and RNA, fibrillation occurs independently of LLPS, resulting in diffuse, thin filamentous networks with minimal evidence of centralized nuclei. This mode of aggregation reflects a fundamentally distinct pathway driven by protein misfolding in the bulk dilute phase. Together, these scenarios illustrate how the timing and interplay of LLPS, gelation, and fibrillation determine the material properties, internal architecture, and morphological diversity of protein condensates. Understanding these kinetic relationships is crucial for deciphering pathological aggregation pathways and may inform strategies for modulating condensate behavior in disease contexts.

**Fig. 7.**
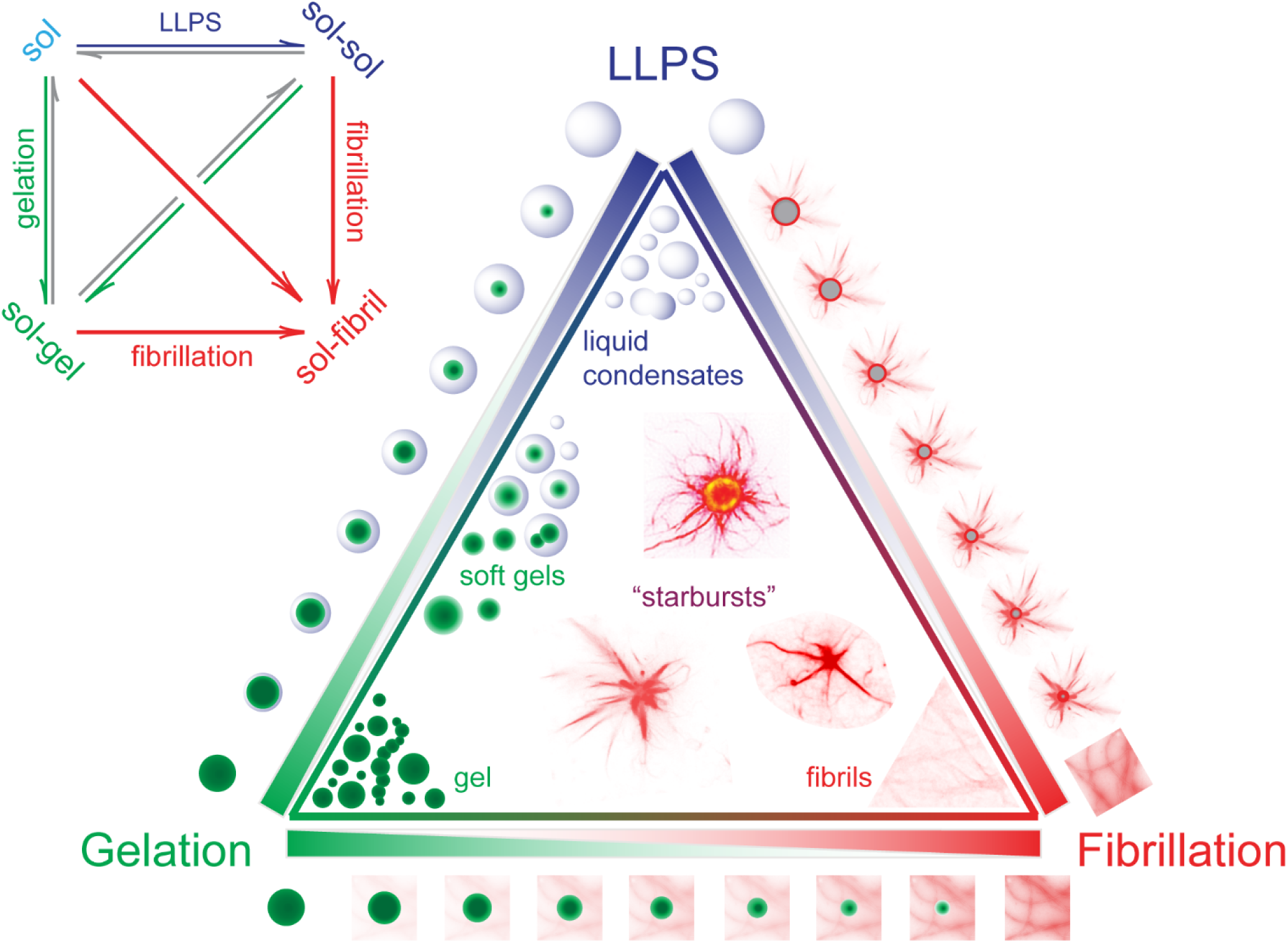
Morphological complexity driven by phase behavior and transitions. The interplay between liquid-liquid phase separation (LLPS), gelation, and fibrillation defines the properties of biomolecular condensates. LLPS mediates transitions between single-phase solutions (sol) and multiphase states (sol-sol). Gelation converts these states into sol-gel networks, while fibrillation emerges from either solution or gel states. For A1PrD, LLPS and gelation processes are largely reversible, whereas fibrillation is irreversible. Depending on the predominant process and the specific phase transitions involved, A1PrD condensates adopt diverse morphologies, including liquid droplets, gels, fibrils, and structurally varied “starburst” aggregates.

Our findings challenge the prevailing assumption that phase-separated compartments invariably correlate with increased protein aggregation. Instead, solution composition significantly influences the balance between pathological and physiological condensates. In our prior studies, we demonstrated that Tubulin actively modulates Tau and α-Synuclein (αSyn) condensates, driving the formation of microtubule-rich, functional condensates, thereby promoting physiological roles (*40*). Extending these observations, our current research highlights a similar protective mechanism involving RNA and A1PrD. Specifically, we demonstrate that RNA presence within A1PrD condensates effectively disrupts or weakens strong homotypic interactions that typically drive protein fibrillation. Thus, RNA-rich conditions, conducive to LLPS, mitigate or significantly delay aggregation, underscoring the protective potential of LLPS in cellular contexts. However, our data also highlight a critical caveat: the aging of these initially protective condensates. As condensates mature, they can undergo gelation or transition into glass-like states, functioning as kinetic traps (*20, 35*). Such arrested condensates effectively immobilize biomolecules indefinitely, reducing protein bioavailability and sequestering essential cellular components. This could explain the pathological hyaline or gel-like inclusions observed for hnRNPA1 and TDP-43 in ALS (*5*). This sustained sequestration can consequently lead to pathological loss-of-function, unless actively resolved by cellular mechanisms, including molecular chaperones or protein quality-control systems(*20*). Together, our findings redefine the understanding of LLPS dynamics, emphasizing the delicate interplay between solution composition and condensate maturation mechanisms that distinguish physiological from pathological outcomes in protein aggregation processes. Insight into these distinct yet interconnected phenomena provides a foundation for therapeutic strategies, enabling targeted interventions aimed at disrupting or modulating pathological aggregation pathways. Ultimately, this offers new opportunities for treating neurodegenerative diseases such as Alzheimer’s disease, Parkinson’s disease, and amyotrophic lateral sclerosis (ALS).

## Acknowledgements

We thank the Baylor College of Medicine (BCM) Optical Imaging & Vital Microscopy Core (OIVM) for use of LSM 880 confocal microscopes. We also thank the Neurological Research Institute (NRI) Microscopy Core for the use of LSM 780 confocal microscopes. We thank Debra Townley (BCM Integrated Microscopy Core) and Lita Duraine (NRI TEM Core) for assistance in TEM data collection. We thank Kim Glass Chu (DesignVictory.us) for the cartoon graphics in the paper. Grant support provided by NINDS, NIH to A.C.M.F (R21 NS107792 and R01 NS105874), and by NIGMS, NIH (R01 GM122763) and the Welch Foundation (Q-2097-20220331) to J.C.F.

## Author contributions

Conceptualization, J.C.F. and A.C.M.F.; Methodology, A.C.M.F., J.C.F. and K.-J.C.; Software and Formal Analysis, M.D.Q., K.-J.C., U.C., S.-C.J.L., P.S.T., C.C.F., J.C.F. and A.C.M.F.; Investigation, M.D.Q., K.-J.C., P.S.T., C.C.F., J.C.F. and A.C.M.F.; Visualization, Writing-Original Draft, Review, & Editing: J.C.F. and A.C.M.F.; Supervision, J.C.F. and A.C.M.F.; Funding Acquisition, J.C.F. and A.C.M.F.

## Data Availability

The authors declare that the main data supporting the findings of this study are available within the article and its supplementary information files. Extra data are available from the corresponding authors upon request.

---

**Fig. S1.**
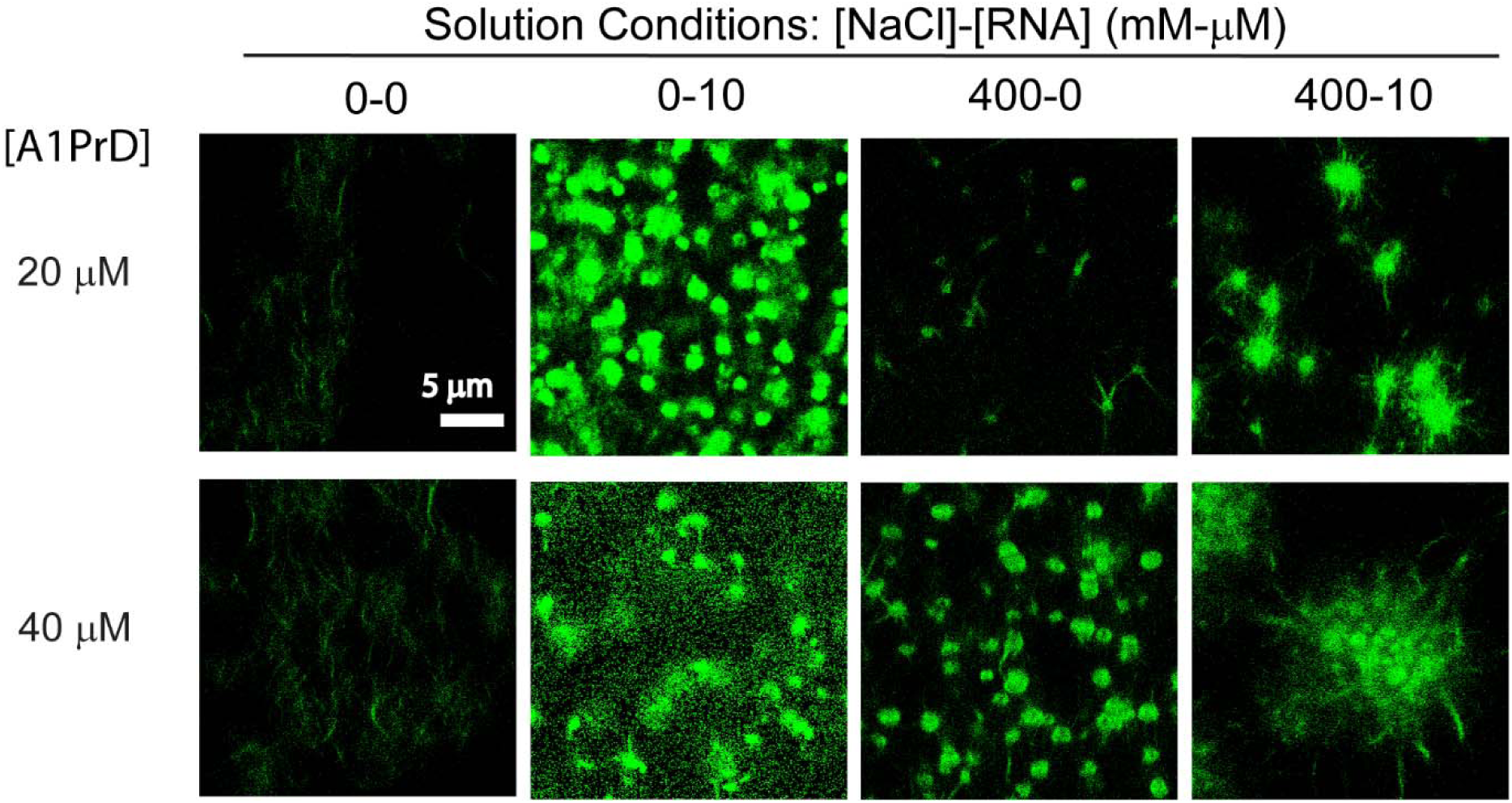
Dependence of condensate and aggregate sizes on A1PrD concentration. Confocal microscopy images of A1PrD (100 nM A1PrD-A488 combined with either 20 or 40 µM unlabeled A1PrD) under varying solution conditions ([NaCl]-[RNA], mM-µM: 0-0, 0-10, 400-0, and 400-10).

**Fig. S2.**
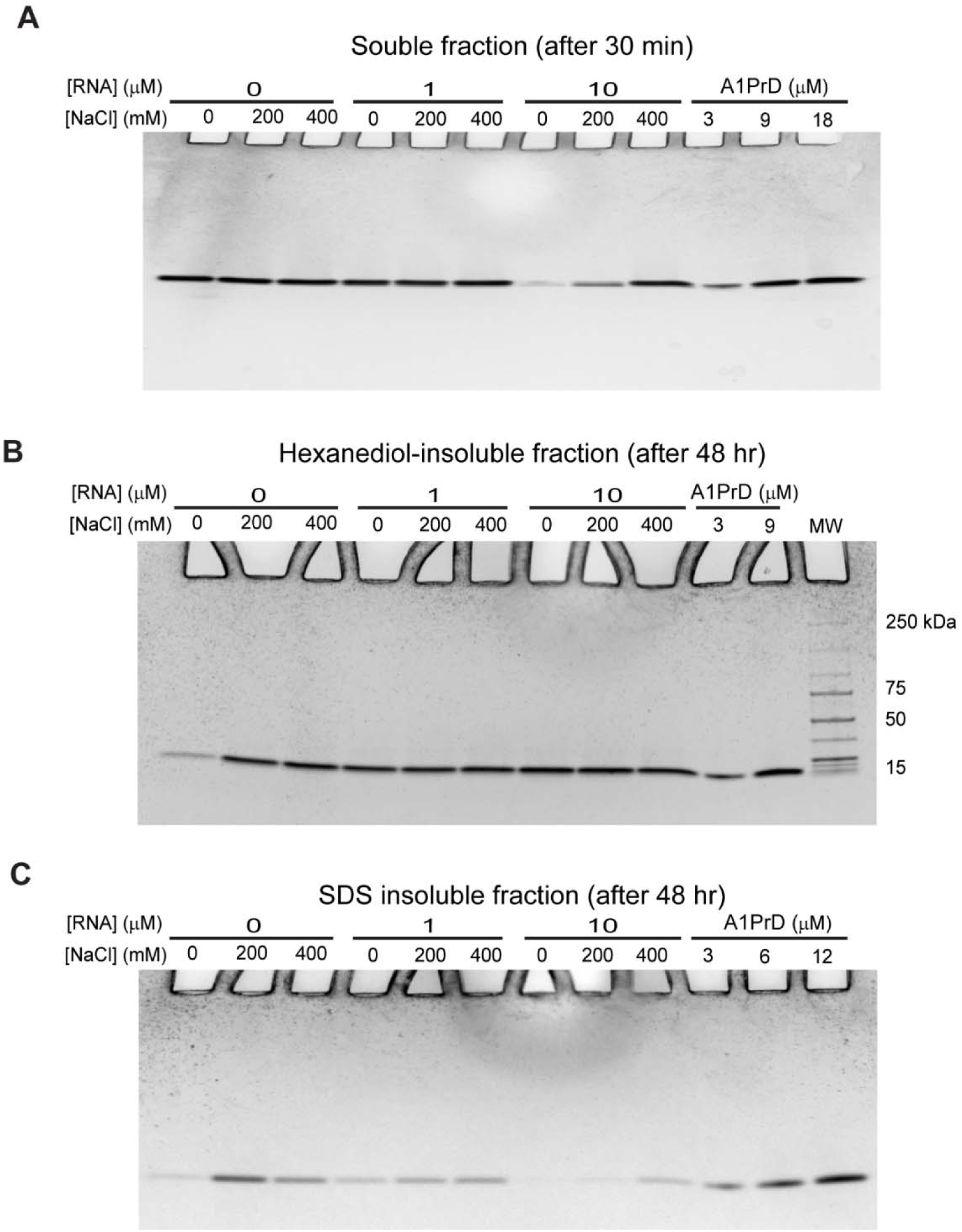
Anti-correlation between A1PrD LLPS potential and SDS-insoluble fibril formation. 20 µM A1PrD samples were prepared under various salt and RNA conditions in αβγ buffer. **(A)** SDS-PAGE analysis of soluble fractions (supernatants collected after 30 min of LLPS initiation). Similar results were confirmed in three independent experiments. **(B)** SDS-PAGE analysis of 1,6-hexanediol-insoluble fractions (pellets collected after 48 hr incubation in 20% 1,6-hexanediol). Similar results were confirmed in three independent experiments. **(C)** SDS-PAGE analysis of SDS-insoluble fractions (pellets collected after 48 hr incubation in 2% SDS). Similar results were confirmed in three independent experiments. Lanes 1-3 represent conditions with 0 µM RNA at increasing salt concentrations (lane 1: 0 mM NaCl; lane 2: 200 mM NaCl; lane 3: 400 mM NaCl). Lanes 4-6 and subsequent lanes represent similar salt gradients with RNA concentrations of 1 or 10 µM. The final three lanes contain 3, 9, and 18 µM A1PrD reference standards for quantification.

**Fig. S3.**
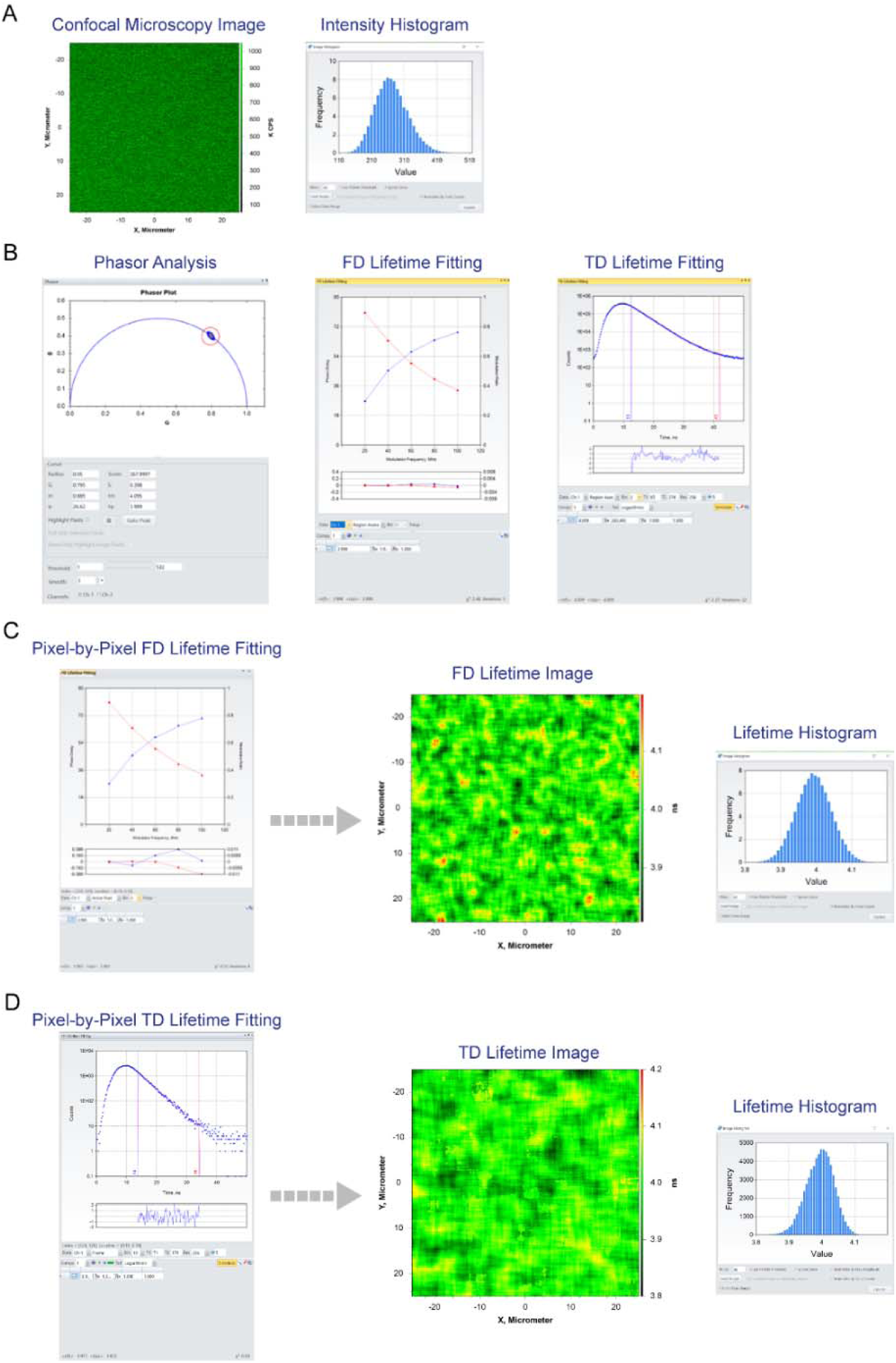
Fluorescence Lifetime Imaging Microscopy (FLIM) data analysis approaches. **(A)** Spatially resolved fluorescence intensity measurements acquired using the ALBA system by x-y scanning, with 10 nM rhodamine 110 in water serving as a control (fluorescence lifetime = 4.0 ns). The histogram distribution of measured intensities confirms the presence of a single molecular species. **(B)** Various methodologies are available for FLIM data analysis, including phasor plot analysis, frequency domain (FD) lifetime fitting, and time domain (TD) lifetime fitting. These analyses can be applied using different formats: pixel-by-pixel, selected groups of pixels (spatially or threshold-based), or globally (using all pixels). **(C-D)** Fluorescence lifetime fitting can be executed pixel-by-pixel across the entire x-y frame, with optional data binning depending on noise levels. Noise is influenced by sample properties and measurement parameters, including dwell time per pixel, frame dimensions, and total pixel count.

**Fig. S4.**
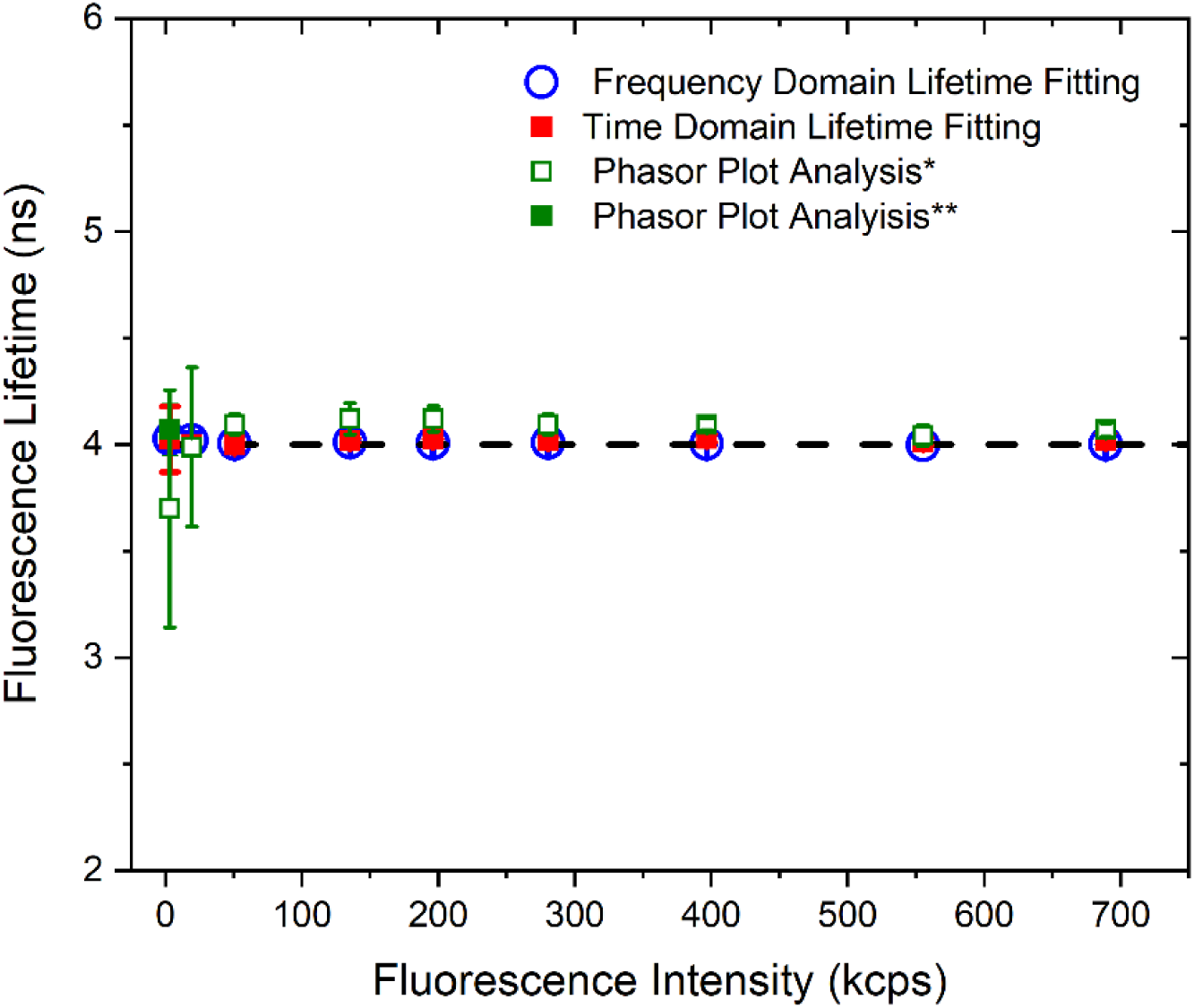
Fluorescence lifetimes measured by FLIM are independent of intensity and analysis method. FLIM data were obtained using 10 nM rhodamine 110 in water as a control, with laser power varying from ∼1-100 µW, corresponding to an average fluorescence intensity range of ∼3-700 kcps. Note: Instrument shutters positioned before the avalanche photodiode detectors close at intensities above ∼800 kcps, and typical experimental settings limit measurements to a maximum intensity of ∼500 kcps to maintain linear detector response. Measured lifetimes remain consistent across different analysis methods and signal intensities, except when signals become extremely low (∼3 kcps), approaching intensities typical of single-molecule measurements (*). Under such low-signal conditions, reliable lifetime measurements can still be achieved by applying increased Gaussian smoothing (**) to the data.

**Fig. S5.**
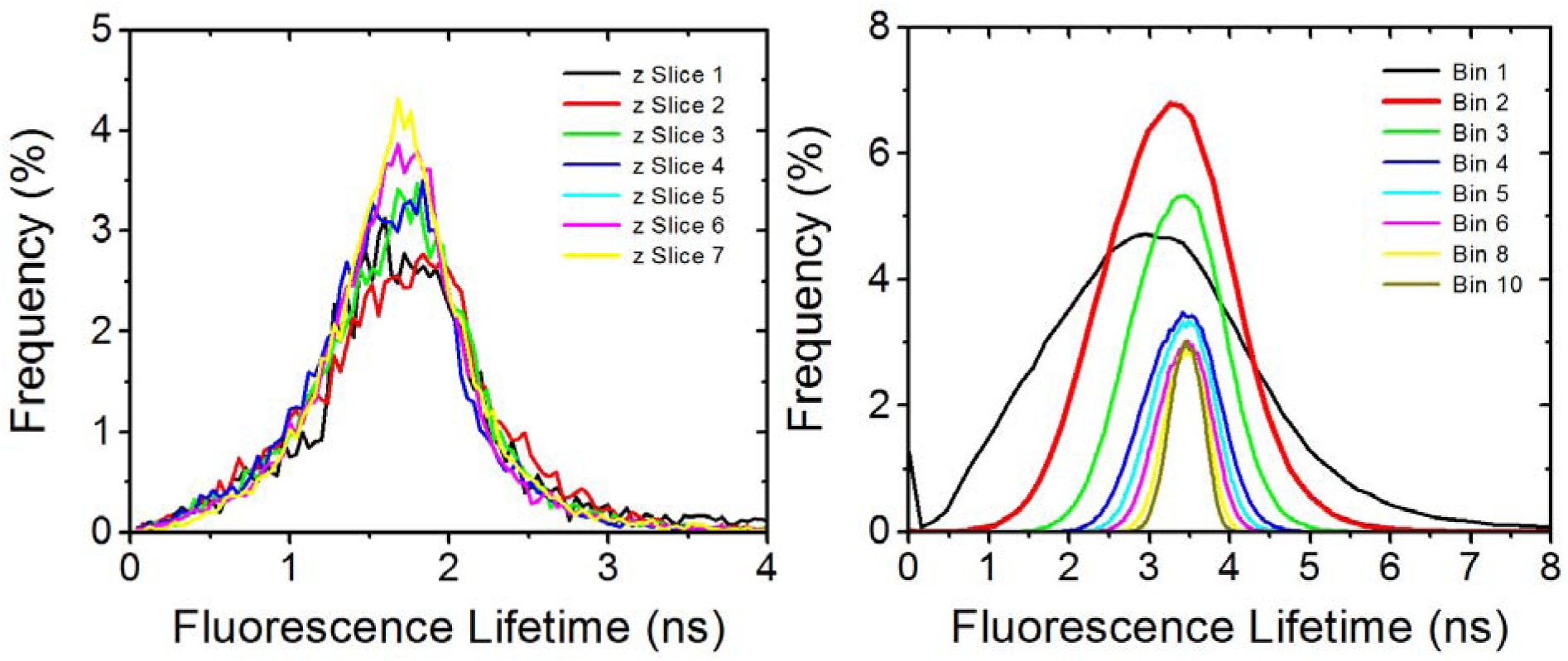
Fluorescence lifetime measurements remain consistent across varying binning times and imaging depths. (Left panel) FLIM lifetime histograms at different binning times. Conditions: ∼70 nM A1PrD-A488 in 10% PEG-8K. (Right panel) FLIM lifetime histograms across various z-slices (each 6 µm thick). Conditions: 20 µM A1PrD with ∼70 nM A1PrD-A488 in 10% PEG-8K.

**Fig. S6.**
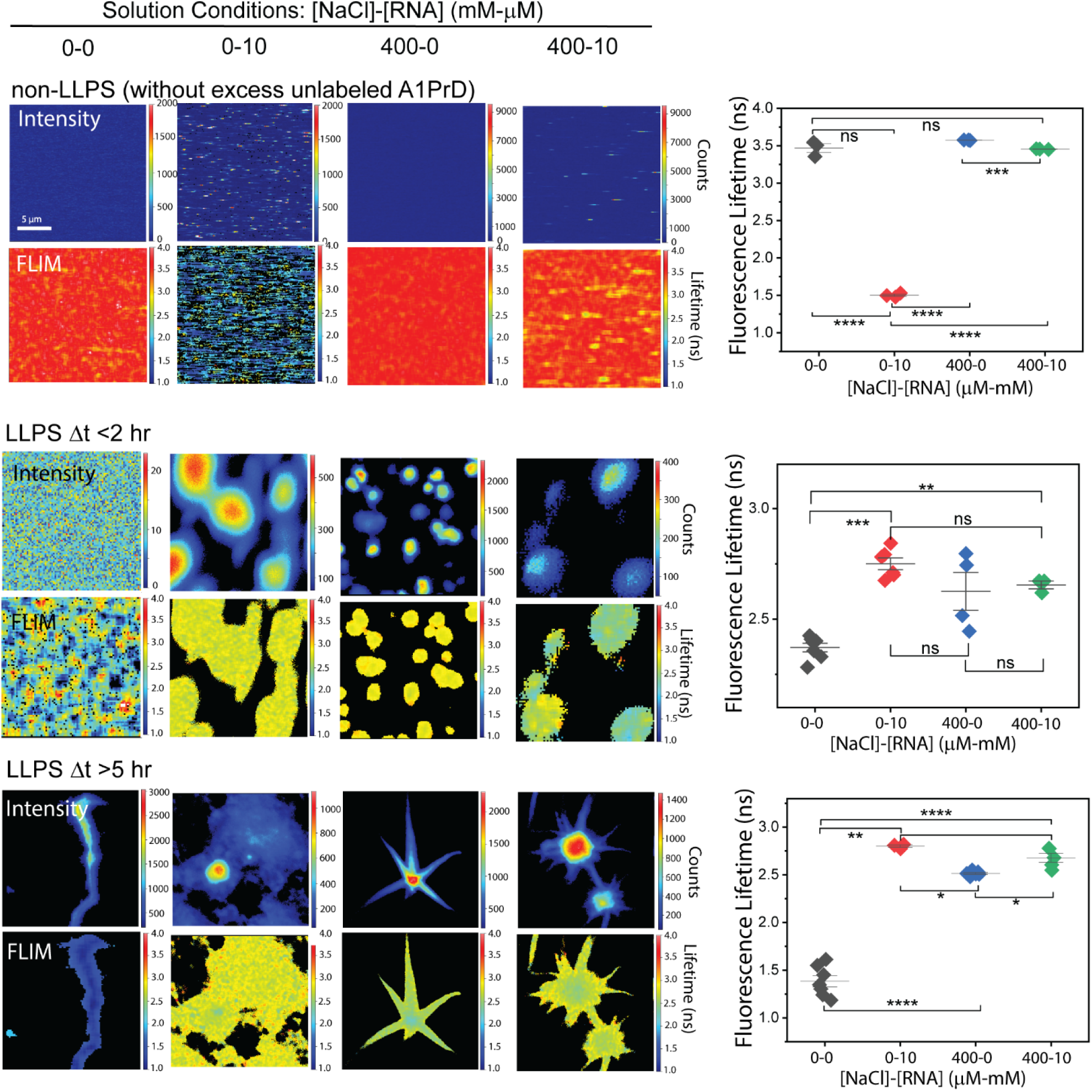
Fluorescence lifetimes of A1PrD under varying solution conditions. (Left, top rows) Confocal fluorescence images illustrating A1PrD starburst maturation at different time points (<2 hr and >5–24 hr post-LLPS initiation). (Left, bottom rows) Corresponding FLIM images. (Right panels) Quantified FLIM lifetimes (mean ± SD; n=3–7 images). Statistical significance determined by two-sided paired Student’s t-tests: ns = not significant, \**P*<0.05, \*\**P*<0.01, \*\*\**P*<0.005, \*\*\*\**P*<0.001.

**Fig. S7.**
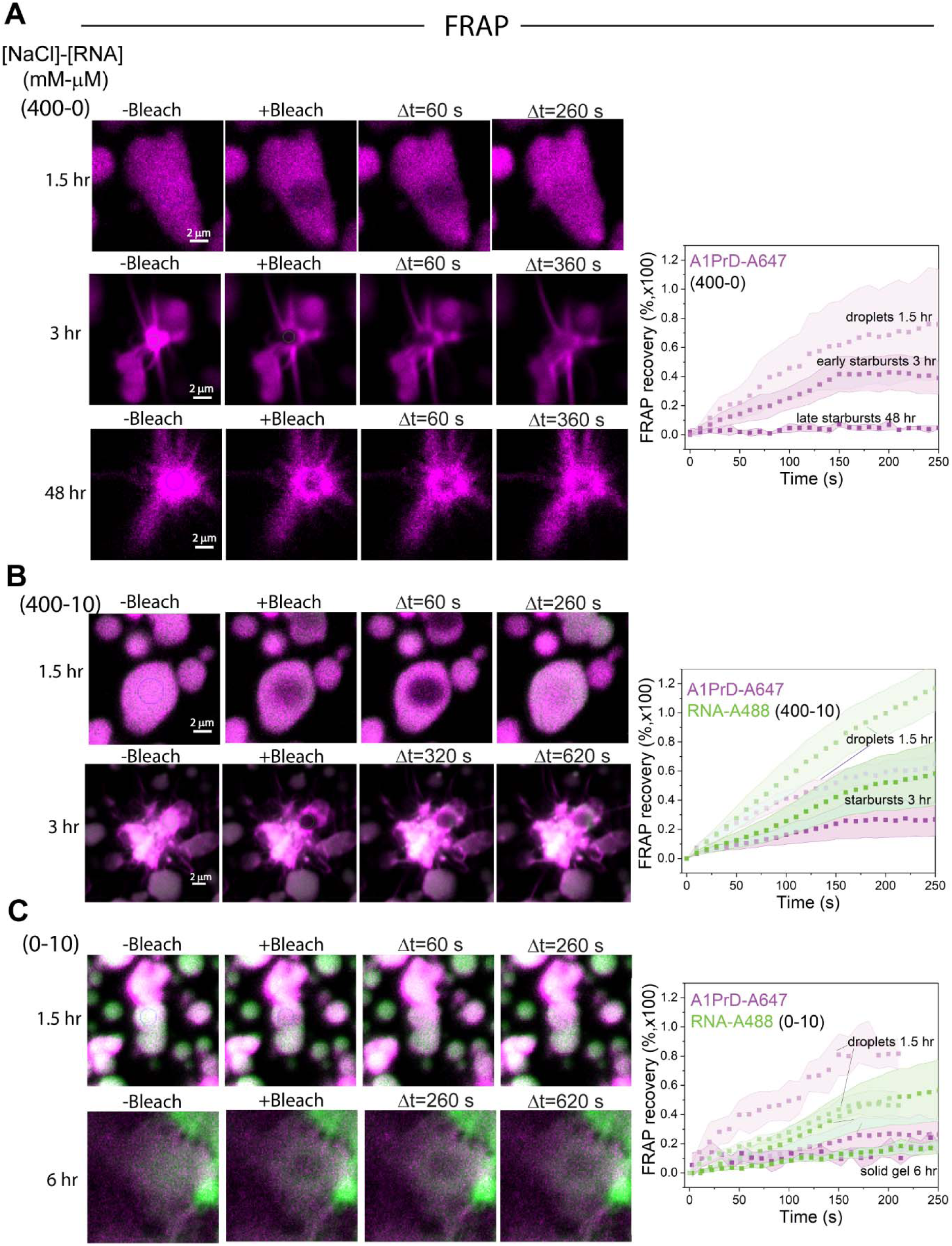
FRAP measurements reveal reduced recovery in aged droplets and starbursts. Representative FRAP images (pre-bleach and post-bleach at specified time intervals) for samples at varying solution conditions ([NaCl]-[RNA], mM-µM: 400-0 **(A)**, 400-10 **(B)**, and 0-10 **(C)**). Corresponding FRAP recovery plots indicate average fluorescence intensities (symbols) and standard deviations (shaded areas) of A1PrD-A647 (purple) and RNA-A488 (green). Sample sizes: 400-0: droplets (n=7), early starbursts (n=3), late starbursts (n=3); 400-10: droplets (n=9), starbursts (n=3); 0-10: droplets (n=4), solid gels (n=5). Darker symbols and shaded areas represent longer-aged samples.

**Fig. S8.**
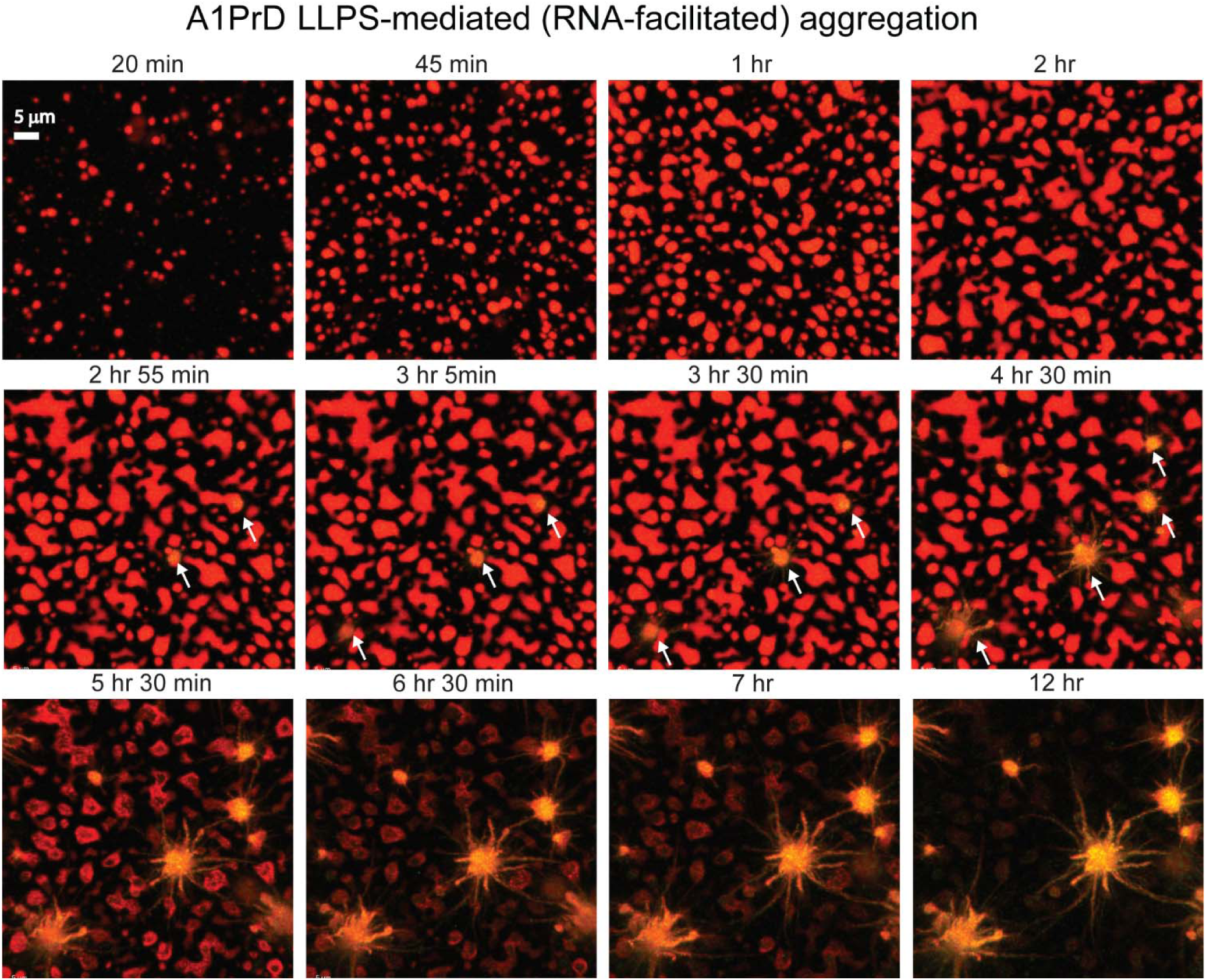
RNA-mediated aging of A1PrD condensates. Time-lapse confocal microscopy images tracking RNA-mediated aging of A1PrD droplets and their subsequent transition to starburst aggregates. Experimental conditions: 20 µM A1PrD, 10 µM RNA, ∼70 nM A1PrD-A647 (red), and 3 µM ThT (green) in αβγ buffer. The average droplet diameter was 2.2 ± 0.4 µm with an aspect ratio (long axis divided by short axis) of 1.3 ± 0.3 (n=414 droplets). Aspect ratio values greater than 1 primarily result from droplet fusion events.

**Fig. S9.**
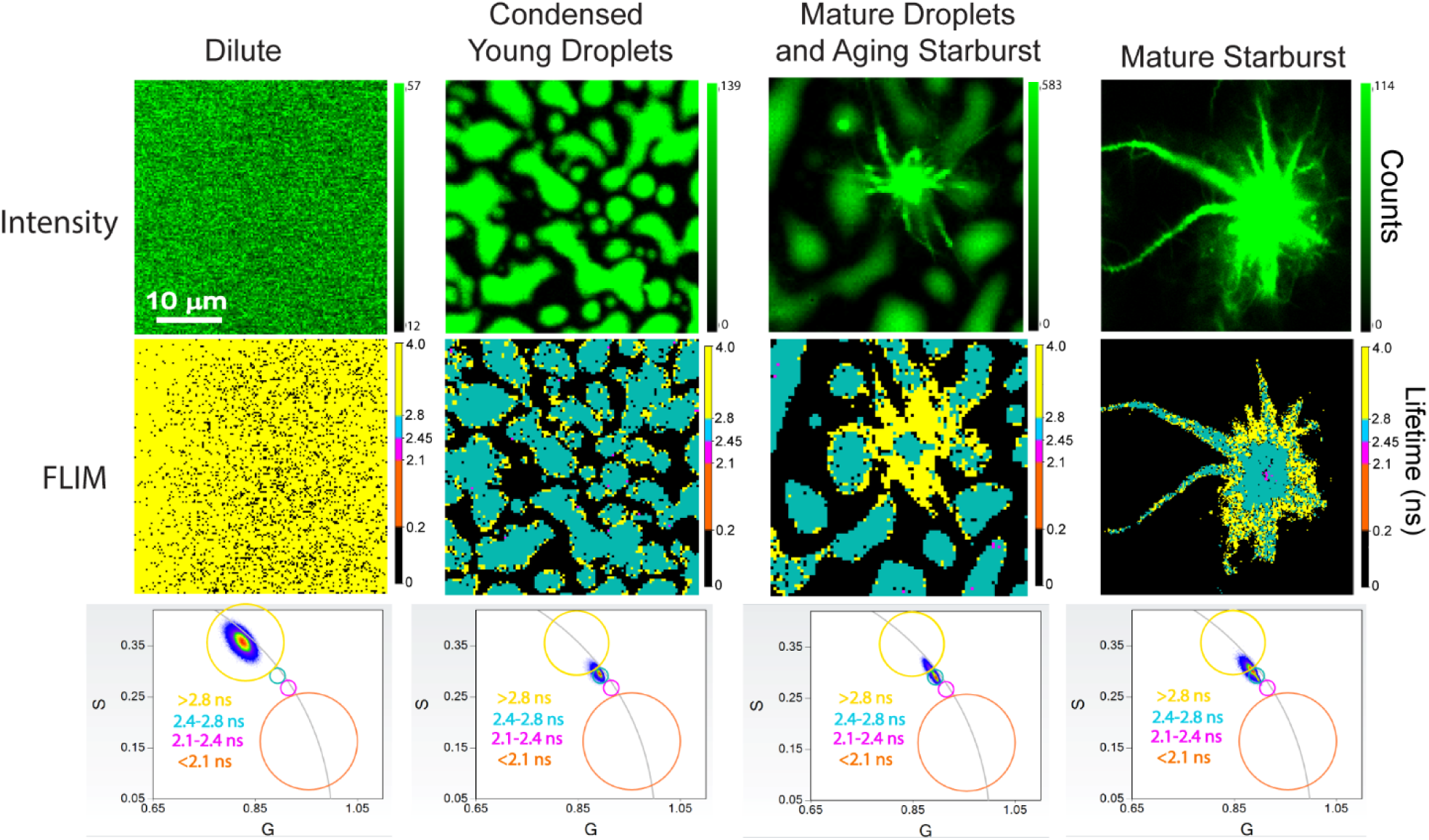
RNA-mediated A1PrD droplets mature into fluid and dynamic starbursts. Confocal microscopy (top row; A1PrD-A488, green) and corresponding FLIM images (middle row) illustrating distinct stages of droplet formation and starburst maturation: non-LLPS dilute conditions, and LLPS conditions at ∼1, ∼4, and ∼22 hr post-LLPS initiation (left to right). Bottom panels show phasor plots with fluorescence lifetimes clustered into four categories: <2.1 ns (orange), 2.1–2.45 ns (magenta), 2.45–2.8 ns (coral blue), and >2.8 ns (yellow). Mean phase lifetimes (τ_ψ_) for each cluster are indicated. Data compiled from 35 measurements across two independent replicates.

**Fig. S10.**
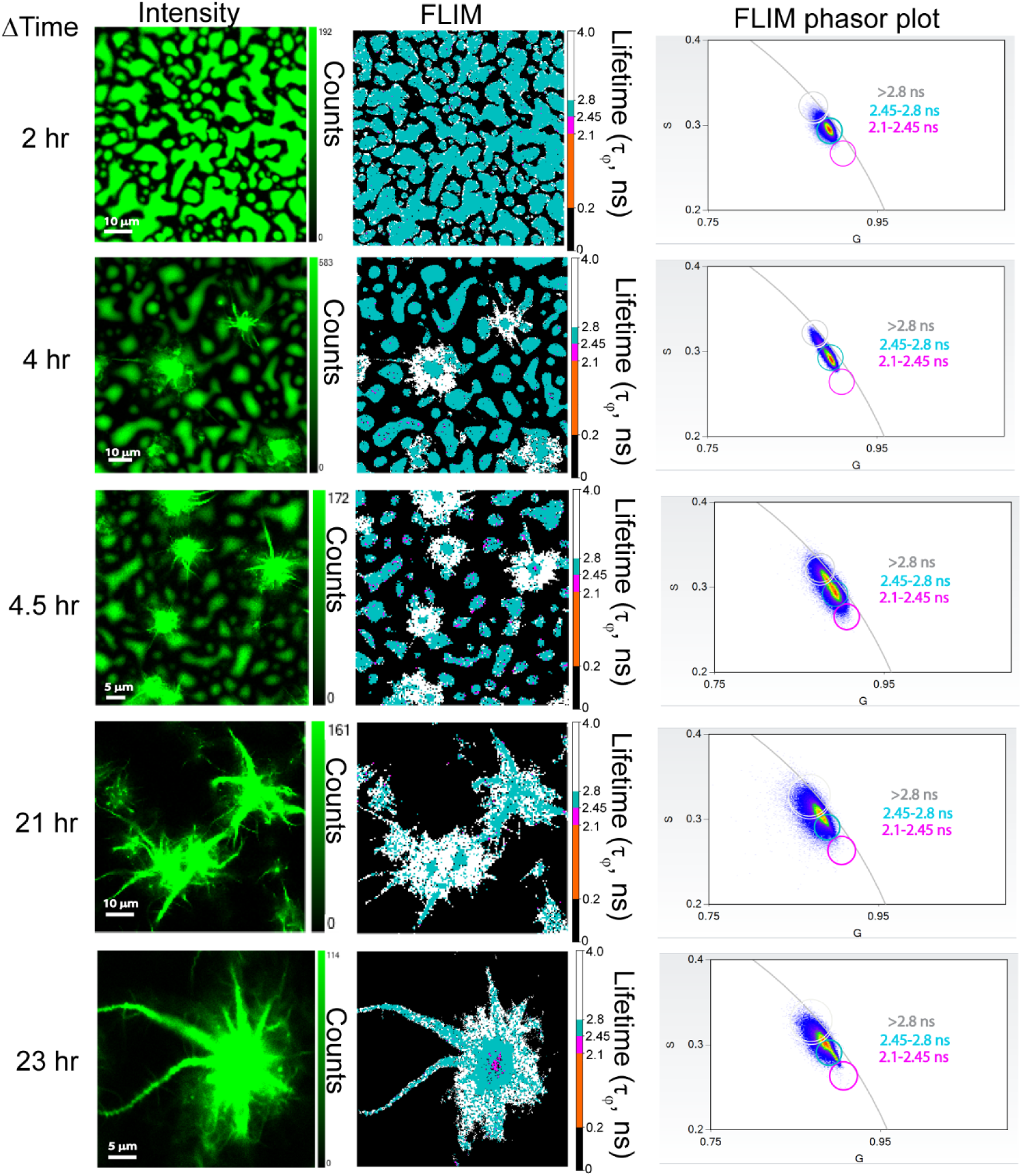
Representative FLIM images of RNA-mediated aging of A1PrD condensates. FLIM experiments performed with 20 µM A1PrD, 10 µM RNA, and ∼70 nM A1PrD-A488 in αβγ buffer. Confocal (left panels) and corresponding FLIM images (middle panels) illustrate progressive aging of RNA-mediated A1PrD droplets into starburst aggregates. Right panels show phasor plots with fluorescence phase lifetimes categorized into three clusters: 2.1–2.45 ns (magenta), 2.45–2.8 ns (coral blue), and >2.8 ns (white/gray).

**Fig. S11.**
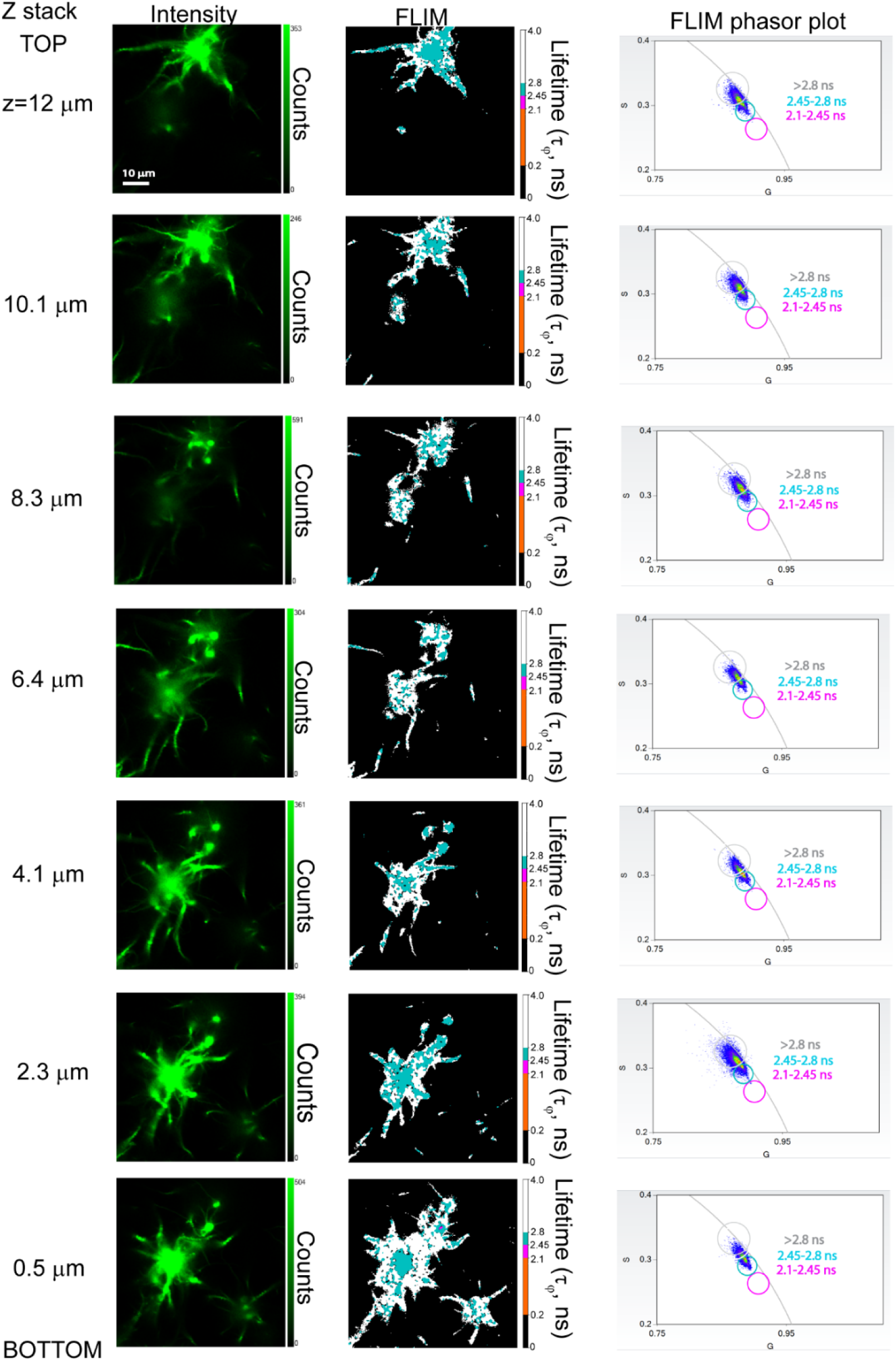
Representative z-stack FLIM images monitoring RNA-mediated aging of A1PrD condensates. Confocal microscopy (left panels) and corresponding FLIM images (middle panels) illustrate A1PrD starburst formation. Phasor plots (right panels) depict fluorescence lifetimes categorized into three clusters: 2.1–2.45 ns (magenta), 2.45–2.8 ns (coral blue), and >2.8 ns (white).

**Fig. S12.**
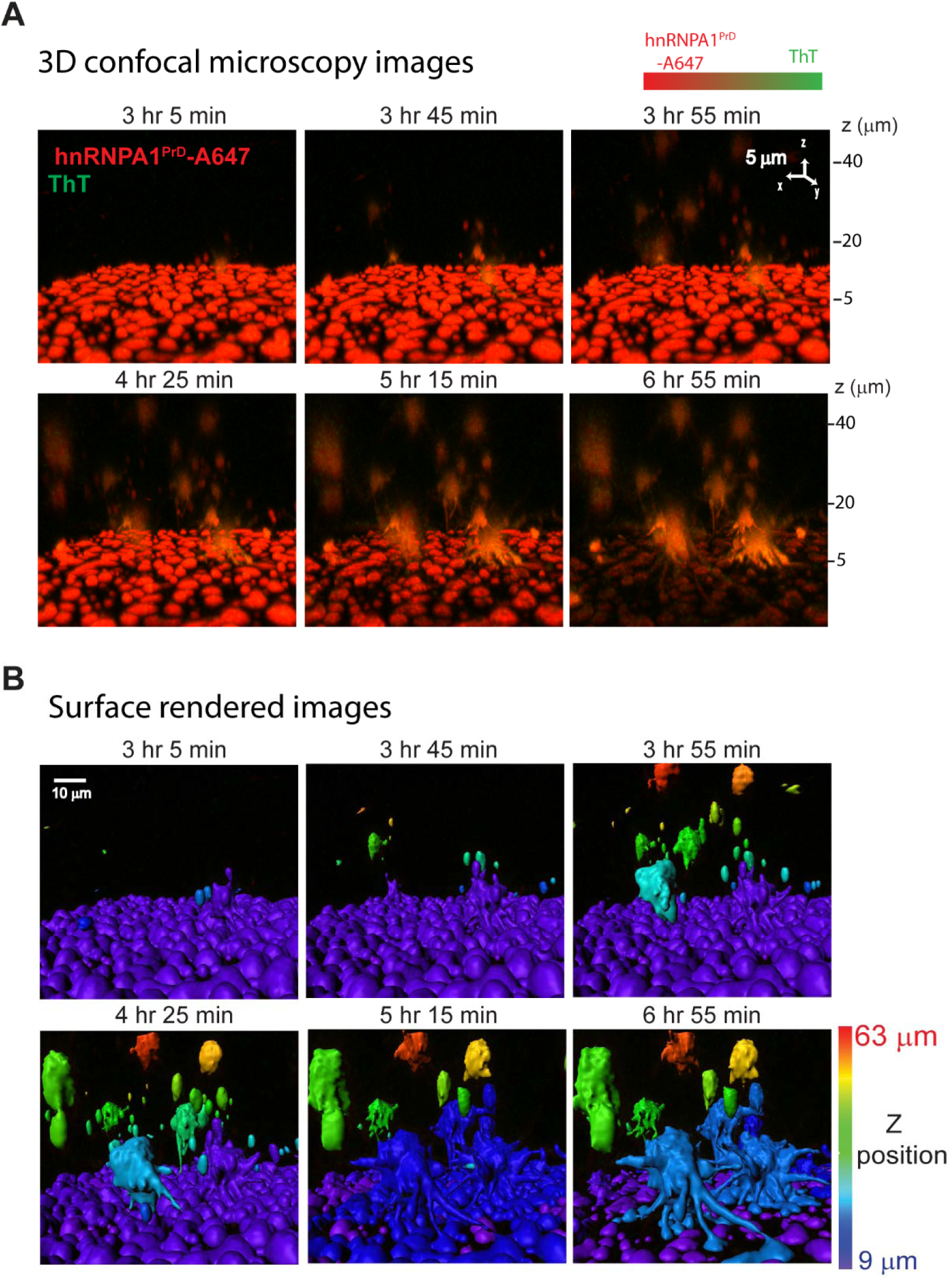
A1PrD starbursts fuse with, siphon material from, and infect young condensates. **(A)** 4D confocal microscopy (time-lapse 3D z-stack) images tracking RNA-mediated aging of A1PrD droplets and subsequent starburst formation. Conditions: 20 µM A1PrD, 10 µM RNA, ∼70 nM A1PrD-A647 (red), and 3 µM ThT (green) in αβγ buffer. **(B)** Surface-rendered representations of images from panel (A), generated using Imaris software.

**Fig. S13.**
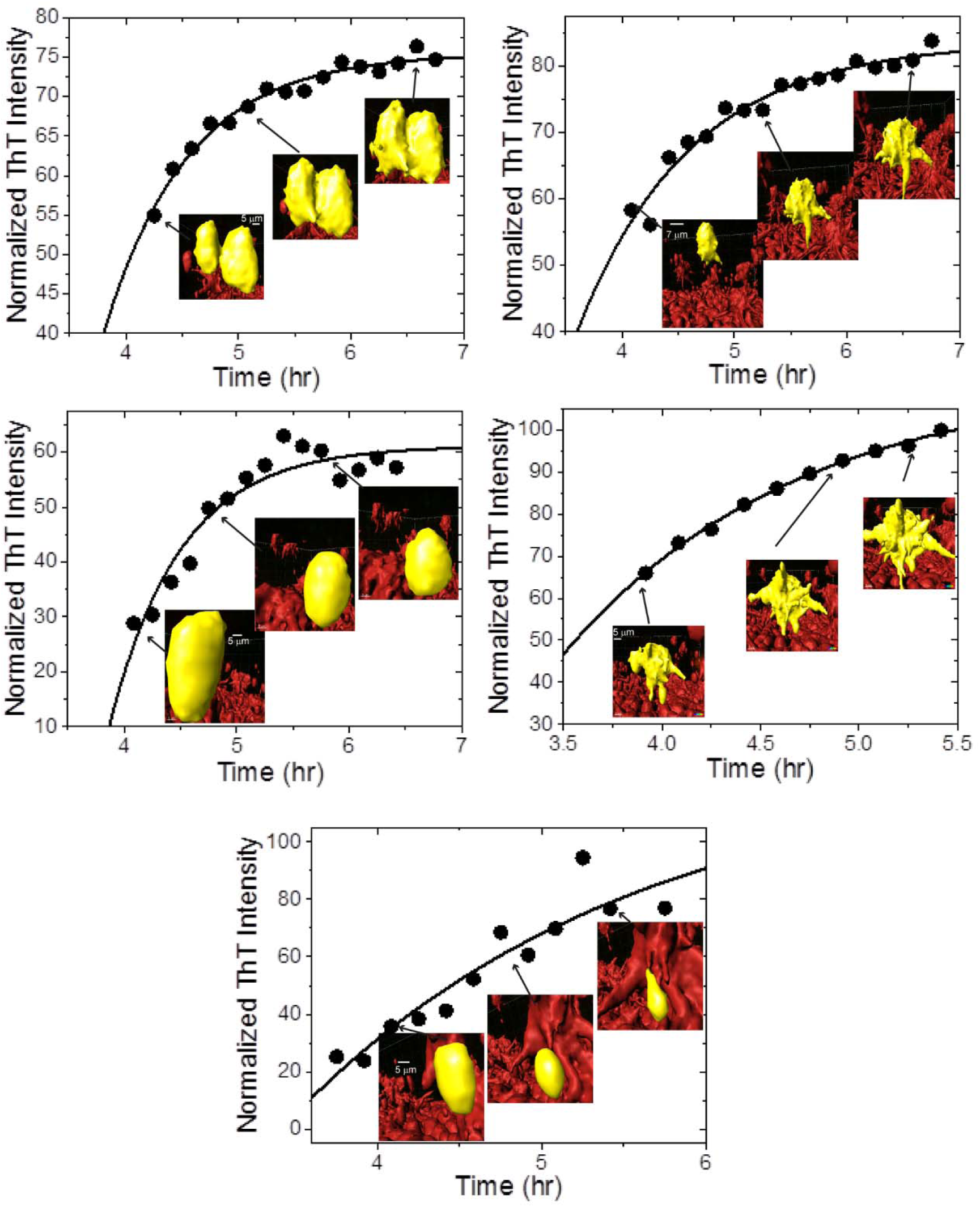
RNA-mediated aging of A1PrD condensates correlates with increased ThT fluorescence. Time-dependent plots of normalized ThT fluorescence intensities (sum of ThT fluorescence intensities divided by condensate surface volumes). Data were fit using a single exponential function (details in Methods).

**Fig. S14.**
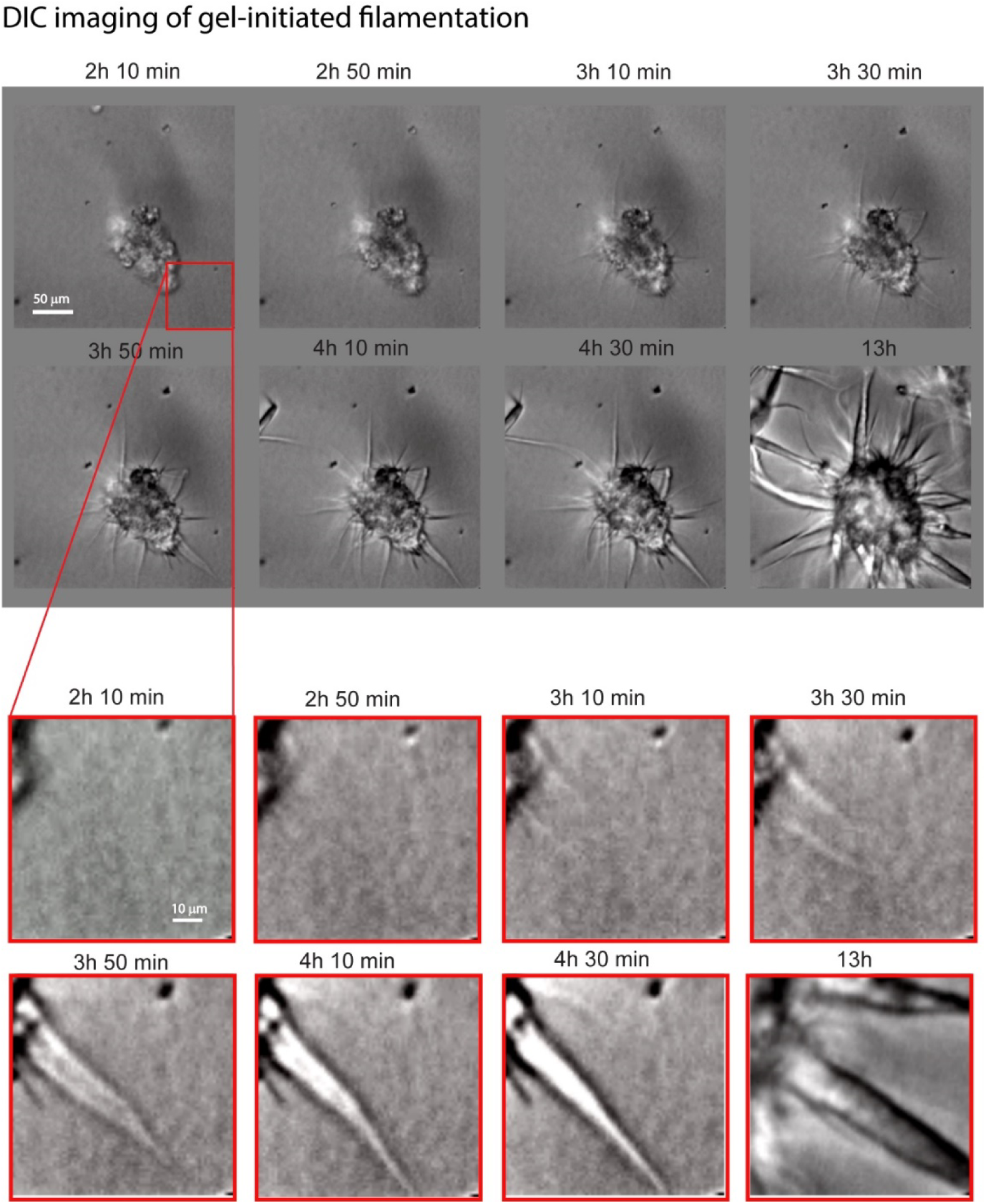
Formation and filamentation of A1PrD starbursts from solid gel clusters. Differential interference contrast (DIC) microscopy images showing different stages of A1PrD gel morphogenesis progressing into starburst aggregates. Conditions: 20 µM unlabeled A1PrD in αβγ buffer (400 mM NaCl, 0 µM RNA).

**Fig. S15.**
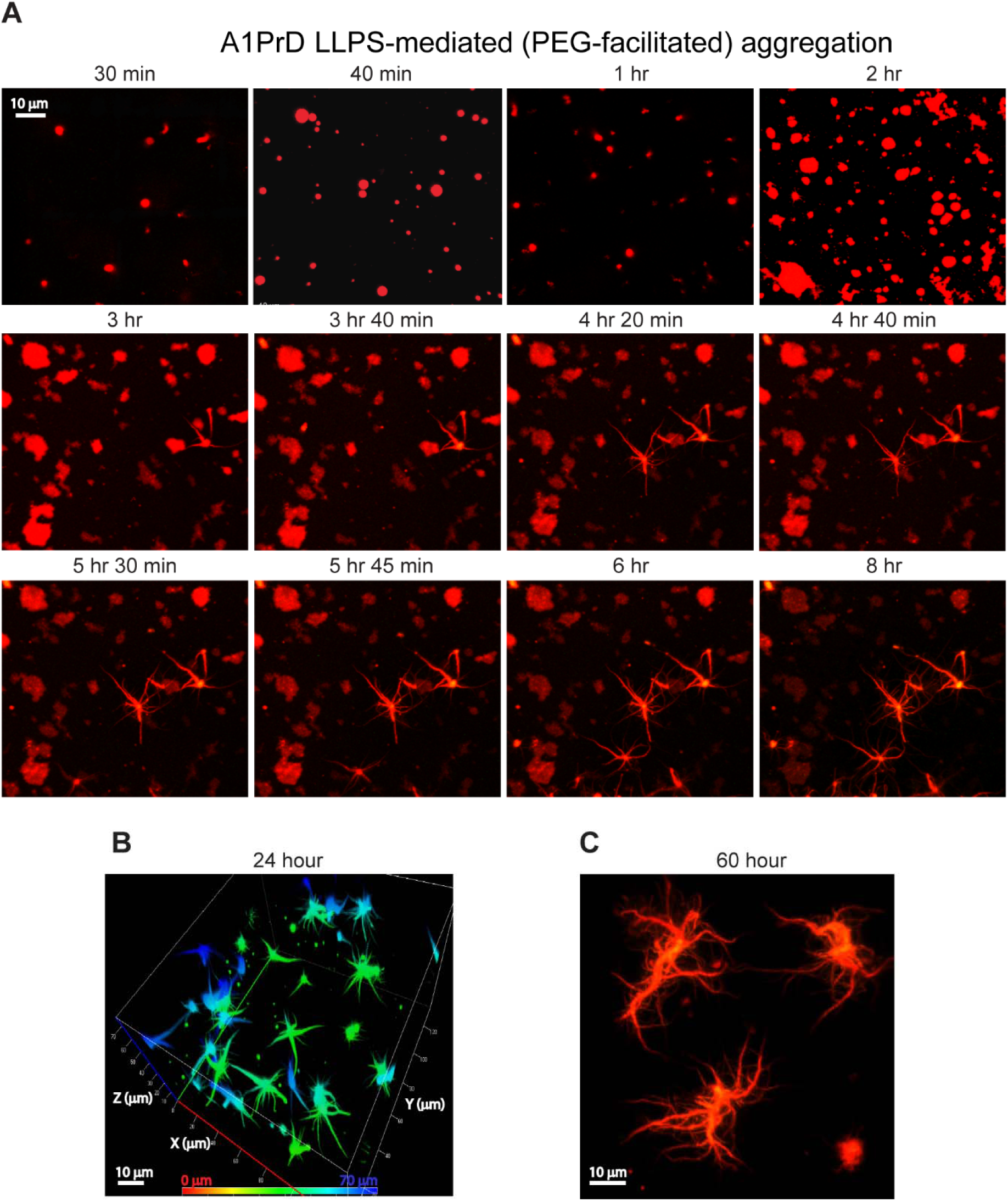
PEG-mediated aging of A1PrD condensates. Crowding-driven protein condensation was examined using 20 µM unlabeled A1PrD, ∼70 nM A1PrD-A647, and 10% w/v PEG-8K in αβγ buffer (with 200 mM NaCl unless noted otherwise). **(A)** Time-lapse confocal microscopy images capturing A1PrD droplet formation, aging, and subsequent starburst development. Droplets formed rapidly (within 30 min), exhibiting spherical morphology with a mean aspect ratio (long axis/short axis) of 1.25 ± 0.2 (n = 65) and diameters averaging 2.4 ± 1.1 µm (n = 65). Larger droplets (>5 µm) frequently demonstrated surface wetting. Images from 20 min to 2 hr represent different regions, whereas subsequent images track the same region. **(B)** 3D confocal image (after 24 hr) highlighting multiple aging starburst structures. **(C)** After 60 hr incubation, mature starbursts exhibited extensive networks of secondary (‘daughter’) spines branching from primary (‘parent’) spines, resembling neurofibrillary tangle morphologies.

**Fig. S16.**
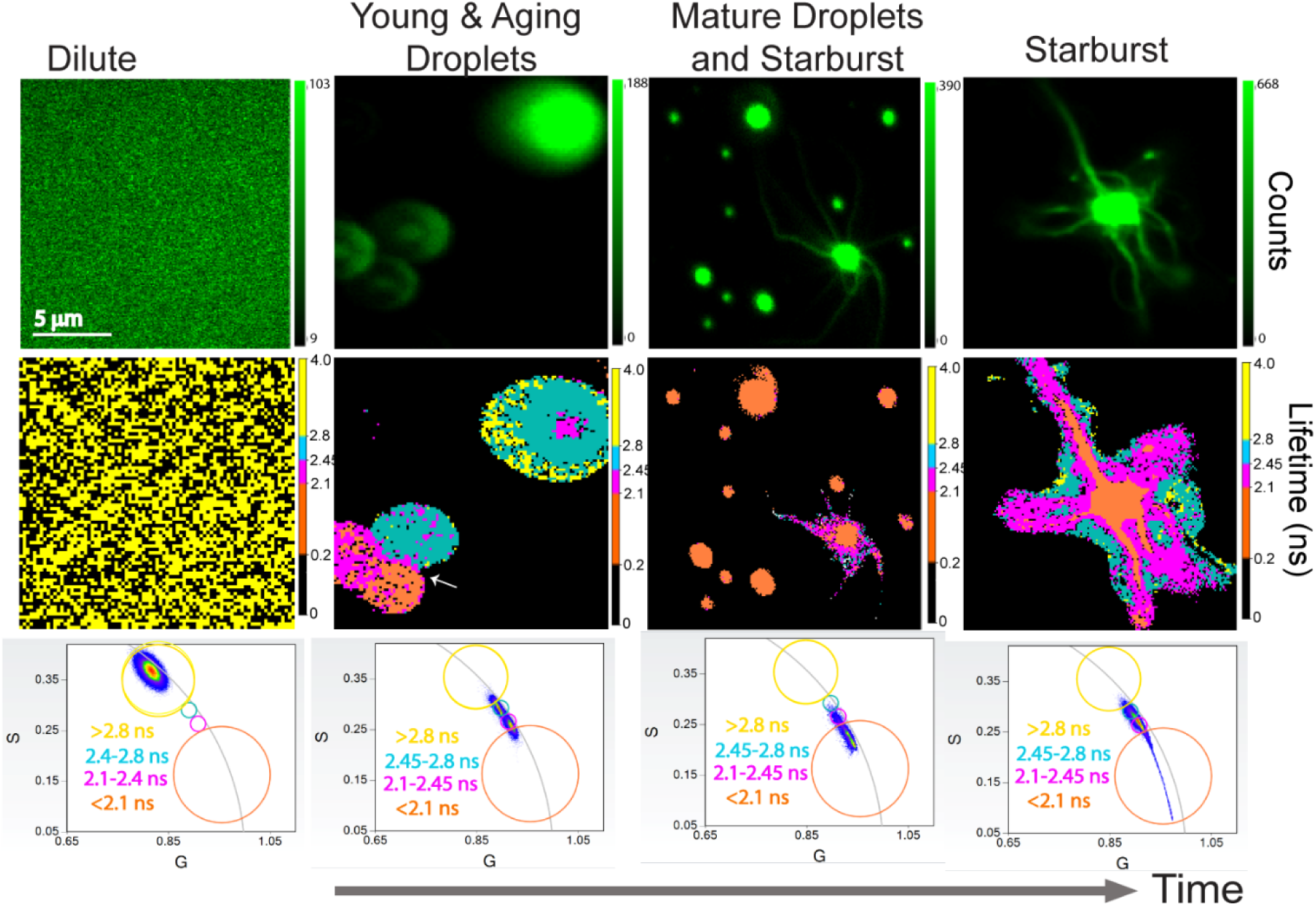
Aging and starburst formation of A1PrD droplets in the presence of PEG. PEG-driven protein condensation monitored using 20 µM unlabeled A1PrD, ∼70 nM A1PrD-A488, and 10% w/v PEG-8K in αβγ buffer (with 200 mM NaCl unless otherwise indicated). Representative confocal microscopy (top row) and FLIM images (middle row) illustrate various stages of droplet maturation and starburst formation: (i) non-LLPS dilute conditions; (ii-iv) LLPS conditions at approximately 5 min, 20 hr, and 24 hr post-LLPS initiation. Corresponding phasor plots (bottom row) depict fluorescence lifetimes grouped into four clusters: <2.1 ns (orange), 2.1–2.45 ns (magenta), 2.45–2.8 ns (coral blue), and >2.8 ns (yellow). Numbers within clusters indicate mean phase lifetimes (τ_ψ_). Data are based on 48 measurements from two independent replicates.

**Fig. S17.**
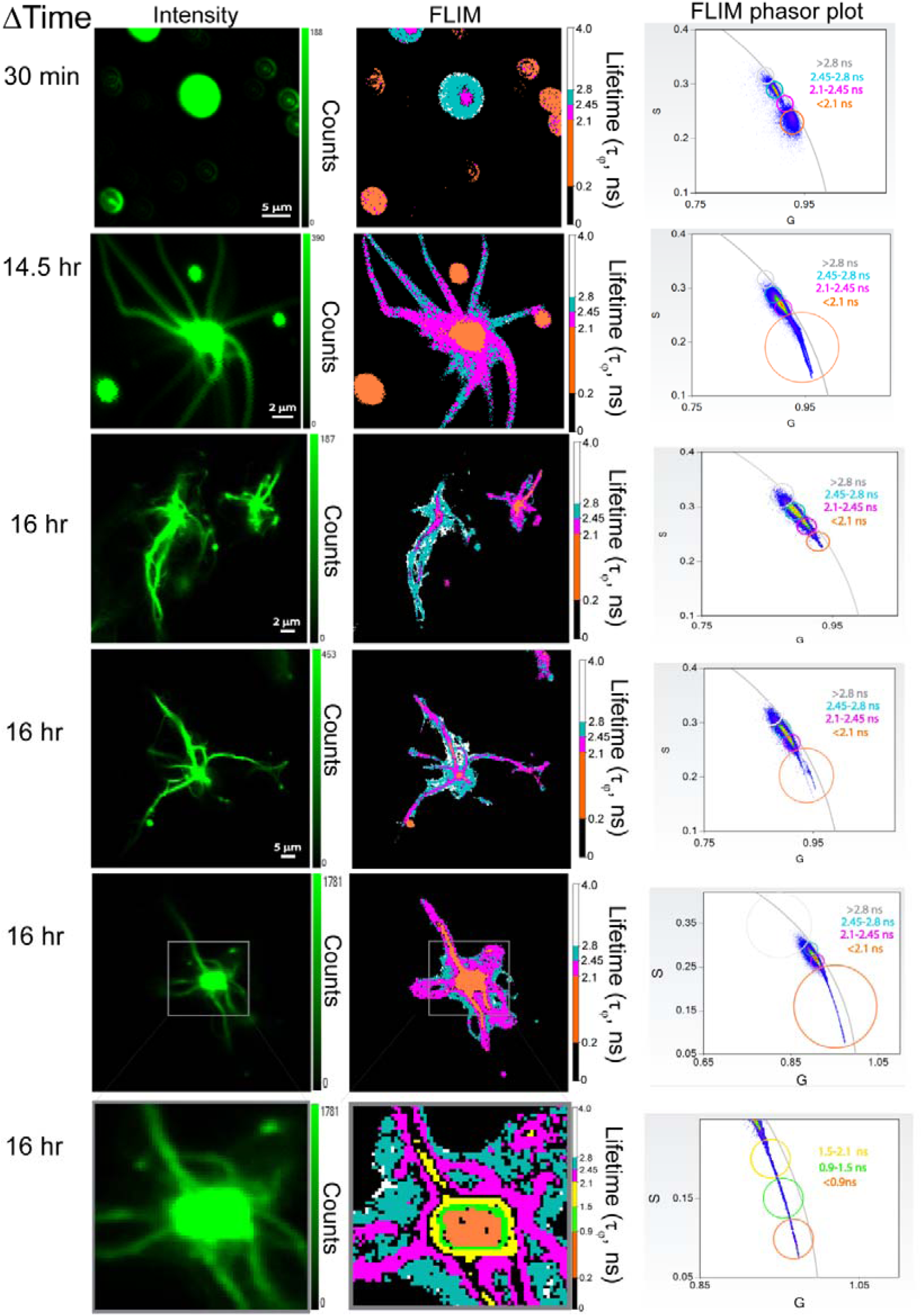
Representative FLIM images tracking PEG-mediated A1PrD droplet aging. FLIM analyses performed on samples containing 20 µM A1PrD, ∼70 nM A1PrD-A488, and 10% w/v PEG-8K in αβγ buffer. Shown are confocal microscopy images (left panels), corresponding FLIM images (middle panels), and phasor plots of FLIM data (right panels), illustrating PEG-driven droplet aging and starburst formation. Fluorescence phase lifetimes (τ_ψ_) are categorized into four clusters: <2.1 ns (orange), 2.1–2.45 ns (magenta), 2.45–2.8 ns (coral blue), and >2.8 ns (grey). The bottom panels are enlarged views of the boxed regions above (16 hr data), highlighting sub-clusters with finer lifetime mapping: <0.9 ns (orange), 0.9–1.5 ns (green), and 1.5–2.1 ns (yellow).

**Fig. S18.**
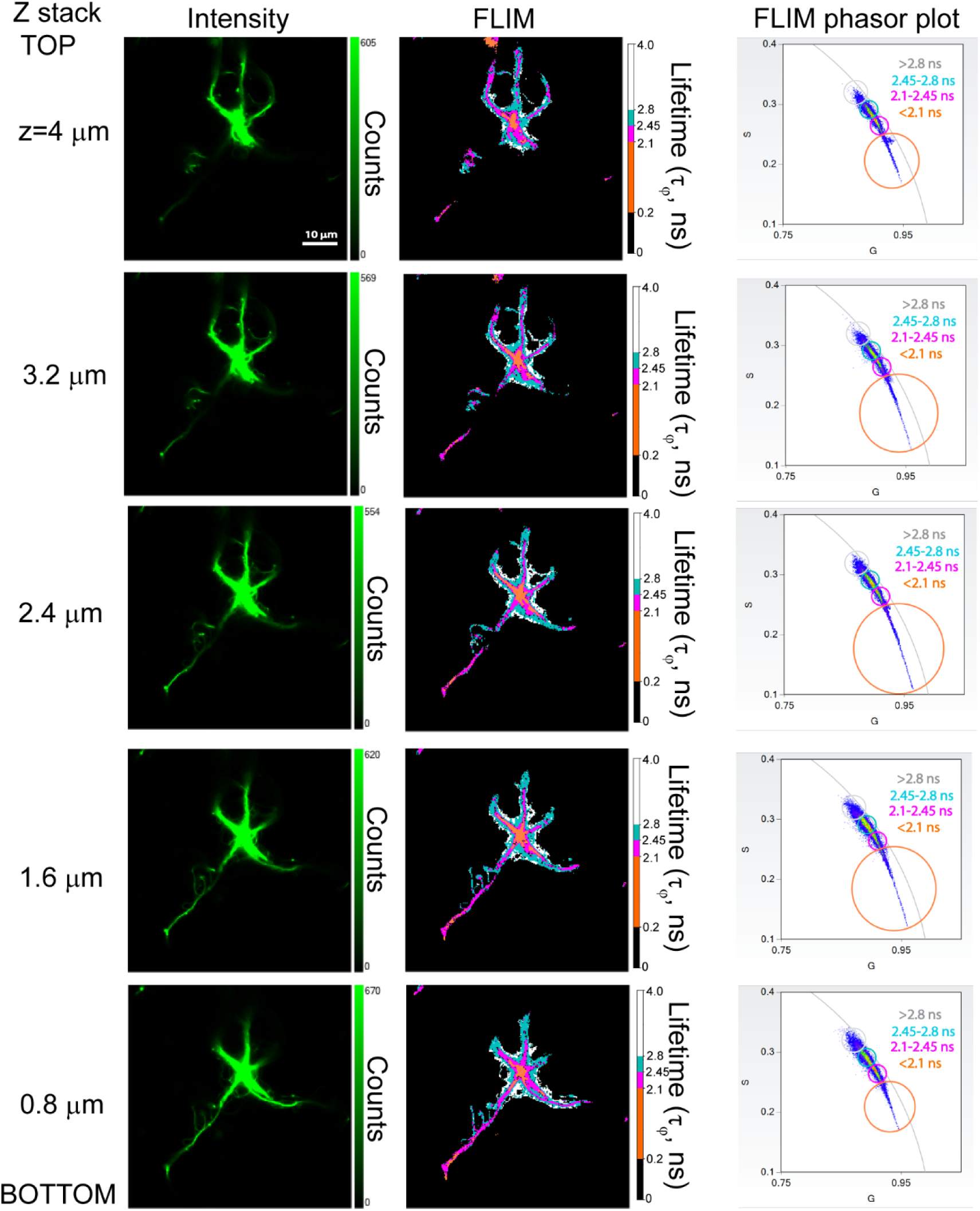
Representative z-stack FLIM images tracking PEG-mediated aging of A1PrD droplets. Confocal microscopy (left panels), corresponding FLIM images (middle panels), and fluorescence lifetime phasor plots (right panels) illustrate PEG-mediated starburst formation from A1PrD droplets. Fluorescence phase lifetimes are categorized into four clusters: <2.1 ns (orange), 2.1–2.45 ns (magenta), 2.45–2.8 ns (coral blue), and >2.8 ns (white/gray).

**Fig. S19.**
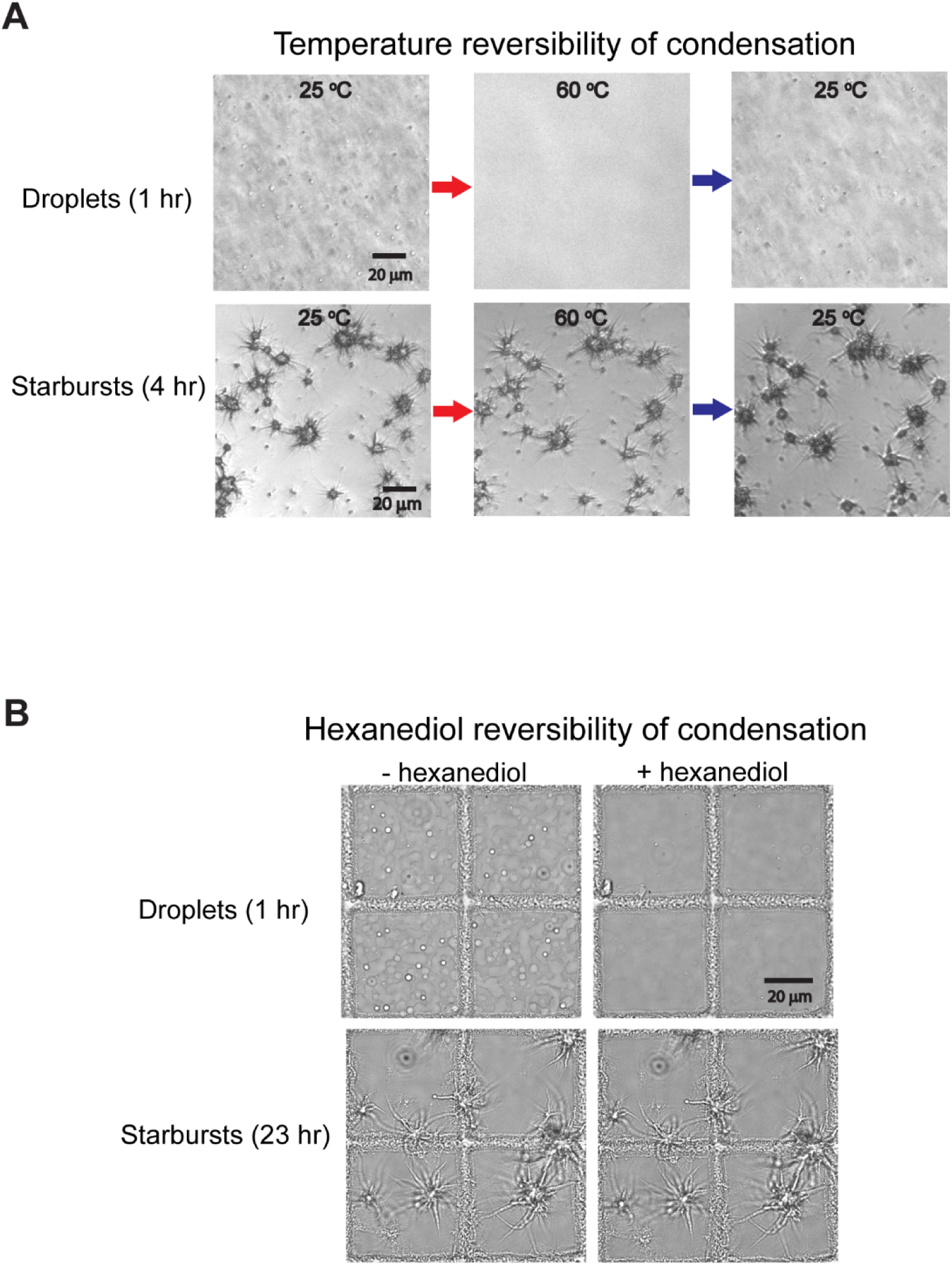
Assessment of droplet and starburst reversibility with temperature and hexanediol. Experiments were conducted using 20 µM A1PrD, 10 µM RNA, and 200 mM NaCl in αβγ buffer. **(A)** DIC microscopy images showing heat-induced dissolution of LLPS droplets, whereas starburst structures remain stable. Results are representative of two independent replicates. **(B)** DIC microscopy images illustrating dissolution of droplets upon addition of 10% 1,6-hexanediol, with starbursts remaining intact. Similar observations were confirmed in two independent replicates.

**Movie S1: SDS-mediated dissolution of A1PrD solid gel condensates.** Samples containing 20 µM unlabeled A1PrD (with 100 nM A1PrD-A488) under conditions of 0 mM NaCl and 10 µM RNA were incubated for 48 hr. Following addition of 1% SDS (final concentration), confocal microscopy images were captured at 30 s intervals over a period of 3.5 min, demonstrating SDS-induced dissolution of most A1PrD gel condensates.

**Movie S2: SDS-mediated dissolution of A1PrD starburst cores.** Samples containing 20 µM unlabeled A1PrD (with 100 nM A1PrD-A488) incubated for 48 hr under conditions of 400 mM NaCl and 0 µM RNA. After addition of 1% SDS (final concentration), confocal microscopy images were captured at 30 s intervals over 3.5 min, demonstrating that SDS treatment effectively dissolves starburst cores but leaves fibrillar extensions intact.

**Movie S3: Aging dynamics of A1PrD RNA-mediated LLPS and aggregation.** Confocal microscopy images capturing an overall view of aged starbursts and residual droplets at 7 hr incubation. Experimental conditions: 20 µM unlabeled A1PrD supplemented with 100 nM A1PrD-A647 and 3 µM ThT, in the presence of 0 mM NaCl and 10 µM RNA.

**Movie S4: Starburst initiation, growth through material sequestration, and condensate infectivity as regulated by fluid-solid balance.** Time-lapse aerial view (xy plane) capturing the aging of A1PrD droplets and their transformation into filamentous starbursts. The movie begins at 115 min post-incubation, displaying combined green (ThT fluorescence) and red (protein fluorescence) signals. Sample conditions are identical to those described in Movie S2.

**Movie S5: Starburst initiation, growth via material sequestration, and condensate infectivity governed by fluid-solid balance.** Time-lapse side view (z-dimension) illustrating the aging of A1PrD droplets and their transition into filamentous starbursts. Sample conditions identical to those in Movie S2.

**Movie S6: Surface-rendered visualization of starburst initiation, growth by material sequestration, and condensate infectivity.** Time-lapse side view (z-dimension) illustrating the aging of A1PrD droplets and their progression into filamentous starbursts. Images are surface-rendered representations derived from Movie S5.

**Movie S7: In-phase fusion, siphoning, and infectivity of A1PrD starbursts.** Time-lapse side view (z-dimension) showing aging and transition of A1PrD droplets into filamentous starbursts. Surface-rendered images represent a different spatial region than Movies S3–S5, with color coding indicating object mass center positions along the z-axis.

**Movie S8: Initiation and growth of A1PrD starbursts without RNA.** 3D time-lapse confocal microscopy images (recorded from 70 min to 4 hr at 10 min intervals) depicting the aging of A1PrD droplets and their transformation into filamentous starbursts. Experimental conditions: 30 µM unlabeled A1PrD (with 100 nM A1PrD-A488), 400 mM NaCl, and 0 µM RNA in αβγ buffer.

**Movie S9: Initiation of filamentous starbursts from A1PrD solid gel clusters.** Time-lapse DIC microscopy (recorded from 70 min to 13 hr at 10 min intervals), aerial view (xy plane), showing the transformation of A1PrD solid gels into filamentous starbursts. Experimental conditions: 60 µM unlabeled A1PrD, 400 mM NaCl, and 0 µM RNA in αβγ buffer.

**Movie S10: PEG-mediated aging of A1PrD droplets drives filament formation from individual condensates transitioning from liquid to solid phases.** Time-lapse fluorescence confocal microscopy recorded from 70 min to 6 hr at 10 min intervals following the initiation of condensation.

**Movie S11: Thermal reversibility of A1PrD liquid droplets.** DIC microscopy time-lapse imaging showing dissolution of A1PrD droplets upon heating to 60°C and subsequent reformation upon cooling to 25°C.

## References

1. M. Goedert, R. Jakes, M. G. Spillantini, The Synucleinopathies: Twenty Years On. J Parkinsons Dis 7, S53–S71 (2017).

2. J. M. Pearce, The Lewy body. J Neurol Neurosurg Psychiatry 71, 214 (2001).

3. C. W. Olanow, D. P. Perl, G. N. DeMartino, K. S. McNaught, Lewy-body formation is an aggresome-related process: a hypothesis. Lancet Neurol 3, 496–503 (2004).

4. S. Zarei et al., A comprehensive review of amyotrophic lateral sclerosis. Surg Neurol Int 6, 171 (2015).

5. H. Mizusawa, Hyaline and Skein-like Inclusions in Amyotrophic Lateral Sclerosis. Neuropathology 13, 201–208 (1993).

6. I. R. Mackenzie, R. Rademakers, M. Neumann, TDP-43 and FUS in amyotrophic lateral sclerosis and frontotemporal dementia. Lancet Neurol 9, 995–1007 (2010).

7. E. E. Connolly, J. F. Ervin, B. L. Plassman, K. A. Welsh-Bohmer, S. J. Wang, Star-shaped TDP-43 inclusions in the oldest-old. J Neuropathol Exp Neurol 84, 356–359 (2025).

8. A. Zbinden, M. Perez-Berlanga, P. De Rossi, M. Polymenidou, Phase Separation and Neurodegenerative Diseases: A Disturbance in the Force. Dev Cell 55, 45–68 (2020).

9. B. Wolozin, P. Ivanov, Stress granules and neurodegeneration. Nat Rev Neurosci 20, 649–666 (2019).

10. S. Boeynaems et al., Protein Phase Separation: A New Phase in Cell Biology. Trends Cell Biol 28, 420–435 (2018).

11. A. Patel et al., A Liquid-to-Solid Phase Transition of the ALS Protein FUS Accelerated by Disease Mutation. Cell 162, 1066–1077 (2015).

12. A. Molliex et al., Phase separation by low complexity domains promotes stress granule assembly and drives pathological fibrillization. Cell 163, 123–133 (2015).

13. Y. R. Li, O. D. King, J. Shorter, A. D. Gitler, Stress granules as crucibles of ALS pathogenesis. J Cell Biol 201, 361–372 (2013).

14. M. Ramaswami, J. P. Taylor, R. Parker, Altered ribostasis: RNA-protein granules in degenerative disorders. Cell 154, 727–736 (2013).

15. Y. Shin et al., Spatiotemporal Control of Intracellular Phase Transitions Using Light-Activated optoDroplets. Cell 168, 159–171 e114 (2017).

16. S. Wegmann et al., Tau protein liquid-liquid phase separation can initiate tau aggregation. EMBO J, (2018).

17. W. M. Babinchak et al., The role of liquid-liquid phase separation in aggregation of the TDP-43 low-complexity domain. J Biol Chem 294, 6306–6317 (2019).

18. T. R. Peskett et al., A Liquid to Solid Phase Transition Underlying Pathological Huntingtin Exon1 Aggregation. Mol Cell 70, 588–601 e586 (2018).

19. W. M. Babinchak, W. K. Surewicz, Liquid-Liquid Phase Separation and Its Mechanistic Role in Pathological Protein Aggregation. J Mol Biol 432, 1910–1925 (2020).

20. S. Alberti, A. A. Hyman, Biomolecular condensates at the nexus of cellular stress, protein aggregation disease and ageing. Nature reviews. Molecular cell biology 22, 196–213 (2021).

21. C. Morelli et al., RNA modulates hnRNPA1A amyloid formation mediated by biomolecular condensates. Nature chemistry 16, 1052–1061 (2024).

22. M. Linsenmeier et al., The interface of condensates of the hnRNPA1 low-complexity domain promotes formation of amyloid fibrils. Nature chemistry 15, 1340–1349 (2023).

23. Y. Sun et al., The nuclear localization sequence mediates hnRNPA1 amyloid fibril formation revealed by cryoEM structure. Nature communications 11, 6349 (2020).

24. P. S. Tsoi, J. C. Ferreon, A. C. M. Ferreon, Initiation of hnRNPA1 Low-Complexity Domain Condensation Monitored by Dynamic Light Scattering. Int J Mol Sci 25, (2024).

25. P. S. Tsoi et al., Electrostatic modulation of hnRNPA1 low-complexity domain liquid-liquid phase separation and aggregation. Protein Sci 30, 1408–1417 (2021).

26. S. Maharana et al., RNA buffers the phase separation behavior of prion-like RNA binding proteins. Science 360, 918–921 (2018).

27. K. J. Choi et al., A Chemical Chaperone Decouples TDP-43 Disordered Domain Phase Separation from Fibrillation. Biochemistry 57, 6822–6826 (2018).

28. K. Joron et al., Fluorescent protein lifetimes report densities and phases of nuclear condensates during embryonic stem-cell differentiation. Nature communications 14, 4885 (2023).

29. M. D. Quan, S. J. Liao, J. C. Ferreon, A. C. M. Ferreon, Fluorescence Lifetime Imaging Microscopy of Biomolecular Condensates. Methods Mol Biol 2563, 135–148 (2023).

30. W. Chen et al., Fluorescence Self-Quenching from Reporter Dyes Informs on the Structural Properties of Amyloid Clusters Formed in Vitro and in Cells. Nano letters 17, 143–149 (2017).

31. J. Klucken, T. F. Outeiro, P. Nguyen, P. J. McLean, B. T. Hyman, Detection of novel intracellular alpha-synuclein oligomeric species by fluorescence lifetime imaging. FASEB J 20, 2050–2057 (2006).

32. W. Becker, Fluorescence lifetime imaging--techniques and applications. J Microsc 247, 119–136 (2012).

33. M. D. Quan, S.-C. J. Liao, J. C. Ferreon, A. C. M. Ferreon, in Phase-Separated Biomolecular Condensates: Methods and Protocols, H.-X. Zhou, J.-H. Spille, P. R. Banerjee, Eds. (Springer US, New York, NY, 2023), pp. 135–148.

34. M. Kar et al., Phase-separating RNA-binding proteins form heterogeneous distributions of clusters in subsaturated solutions. Proc Natl Acad Sci U S A 119, e2202222119 (2022).

35. L. Jawerth et al., Protein condensates as aging Maxwell fluids. Science 370, 1317–1323 (2020).

36. C. M. Dobson, Protein folding and misfolding. Nature 426, 884–890 (2003).

37. P. G. Vekilov, Metastable mesoscopic phases in concentrated protein solutions. Ann N Y Acad Sci 1161, 377–386 (2009).

38. T. J. Nott et al., Phase transition of a disordered nuage protein generates environmentally responsive membraneless organelles. Mol Cell 57, 936–947 (2015).

39. R. Sharma et al., Liquid condensation of reprogramming factor KLF4 with DNA provides a mechanism for chromatin organization. Nature communications 12, 5579 (2021).

40. L. Lucas, et al., Tubulin transforms Tau and alpha-synuclein condensates from pathological to physiological. bioRxiv, (2025).

