## Supplementary methods for "Modulation of biomolecular aggregate morphology and condensate infectivity"

**Cloning.** The C-terminal hnRNPA1 sequence (residues 186-320; A1PrD) was PCR-amplified from pET9d-hnRNPA1 (Addgene plasmid #23026, gift from Douglas Black) to generate A1PrD (Prion-like Domain of hnRNPA1) plasmid constructs. The primers used in this study are: 5’-ctgtattttcagggacatatggctagtgcttcatccagcc-3’ and 5’-ctcgagtgcggccgcaccttaaaatcttctgccactgcc-3’. Purified PCR product and NdeI/HindIII-digested pET29-mCherry-TEV vector (*developed in-house*) were used for Gibson assembly(*1*). The reactions consisted of 4 μL 2X Gibson Mixture (New England Biolabs) and 4 μL insert-linearized vector mixtures (7-fold molar excess of insert), and were incubated at 50^°^C for 1 hr. The complete mixtures were used to transform chemically competent DH5α cells. The plasmid (pET28-mCherry-A1PrD) containing the insert was verified by DNA sequencing.

**Protein expression and purification.** The pET28-mCherry-A1PrD plasmid was transformed into *E.coli* Rosetta (DE3) competent cells (Novagen) and grown at 37.0°C in Terrific Broth medium containing kanamycin. Cells were induced for expression with 1 mM IPTG at OD_600_ 1.0-1.5. Growth was allowed to continue overnight at 18.0°C, and cells were subsequently harvested by centrifugation. The recombinant fusion protein, N-terminal 6XHis-mCherry-TEV cleavage site-A1PrD, was expressed as a soluble protein (MW 45.5 kDa). For A1PrD purification, cells were suspended in purification buffer (300 mM NaCl, 50 mM Tris, pH 7.5) containing a protease inhibitor cocktail (GenDEPOT), and lysed using a homogenizer (Avestin). Purifications were then carried out either by using a batch/gravity method wherein clarified lysates were applied to HisPur cobalt resin (ThermoFisher Scientific) or by FPLC using Talon columns (GE Healthcare). After extensive washing, 6XHis-tagged protein was eluted using 200 mM imidazole, dialyzed against purification buffer, and cleaved with TEV protease (*prepared in-house*; ~1:50 mass ratio) to remove 6XHis-mCherry tags. The cleaved A1PrD protein was then separated from the 6XHis-mCherry tags by passing the sample through the cobalt resin again. A1PrD fractions were further purified by reverse-phase HPLC (20-60% ACN gradient with 0.1%TFA) using semi-preparative C3 or C18 columns (Agilent).

**Protein fluorescence labeling.** Purified A1PrD were dye-labeled using NHS ester chemistries with Alexa Fluor 488 (A488) or Alexa Fluor 647 (A647) NHS (with 1:10 protein:dye labeling ratio, incubated at RT for 1-3 hr; Thermo Fisher Scientific), similar to previously described methods(*2, 3*). The labeled samples were subsequently purified by reverse-phase HPLC (20-60% ACN gradient with 0.1%TFA), and lyophilized and stored at -80°C until further use.

**Fluorescence microscopy imaging.** Confocal fluorescence imaging was performed using LSM710 or LSM880 confocal microscopes (Zeiss) for timelapse z-stack experiments. DIC and epifluorescence imaging were performed using a Nikon Ti2 SE microscope (Nikon) and EVOS FL Imaging System (Thermo Fisher Scientific). For microscopy phase diagrams varying NaC] and RNA concentrations, fluorescence imaging was performed using a spinning disk Nikon Ti2 SE microscope (Nikon) equipped with a CSU-W1 confocal scanner unit (Yokogawa).

*Sample preparation for fluorescence imaging.* For PEG- or RNA-mediated LLPS experiments, separate stock solutions of unlabeled (880 μM) and labeled A1PrD (29 μM; either A647- or A488-labeled) were prepared and dissolved in 4 M urea in 200 mM NaCl, αβγ buffer (10 mM acetate, 10 mM phosphate, 10 mM glycine, pH 7.5). Substock solutions of protein mix (1:300 labeled vs. unlabeled) were prepared and frozen in 1-2 μL aliquots at -80°C prior to imaging. Each aliquot was diluted to 20-40 μL volume with other experimental buffer components, generating the final solution conditions for A1PrD (~1:300 labeled to unlabeled ratio; 67 nM labeled:20 μM unlabeled A1PrD) in 100 mM urea, 200 mM NaCl, αβγ buffer, either with 10 μM non-specific RNA (5’-GGGCCCCCGGGUACCGAGCUGCUAAUCAAAACAAAACAAAAGCU-3’; described in Molliex *et al*(*4*) ) or 10% w/v PEG-8K (Hampton Research). For some samples, 3 μM thioflavin T (ThT) were added to monitor amyloid formation.

For experiments in different NaCl and RNA conditions, lyophilized A1PrD protein was dissolved in water and subsequently mixed with stock αβγ buffer containing various NaCl and RNA concentrations to generate the final solution conditions ([NaCl]: 0, 200, and 400 mM; and [RNA]: 0, 1, and 10 µM); pH was individually adjusted for each stock buffer. In some cases, either A647-labeled or A488-labeled A1PrD were added at 1:100 or 1:200 labeled to unlabeled ratio (~100 or 200 nM labeled:20 μM unlabeled A1PrD), or A488-RNA (~50 nM) was added to 10 µM unlabeled RNA (1:200 labeled to unlabeled ratio). Different protein concentrations used in experiments are indicated in the figure legends.

Unless specified otherwise, samples were deposited onto a glass-bottomed microscope dish (MatTek dish, or Ibidi μ-Dish, 35 mm, with or without grid). To minimize evaporation effects in experiments performed at RT, several 20 μL buffer droplets were added around the dish periphery, and dishes were sealed with parafilm. For thermal experiments, samples were covered in paraffin oil or a cover slip (with a diameter bigger than the dish bottom coverslip leaving enough height (~1 mm) for molecular movement within the sample), which was then sealed with nail polish.

*4D (timelapse z-stack) confocal imaging.* Timelapse z-stack images were collected spanning 60-100 μm above the glass surface, and from 10 min to 9 hr of collection at varying time intervals (0 to 10 min per z-stack). For the timelapse z-stack experiment at 55°C, 100 μL sample of A647-labeled or A488-labeled A1PrD (1:300 labeled to unlabeled ratio; 67 nM labeled:20 μM unlabeled A1PrD) with 10 μM RNA in 200 mM NaCl, αβγ buffer were pipetted onto 35 mm (No. 1.5 coverslip) MatTek dishes (MatTeK Corporation) covered with Coverwell^TM^ chambers (Grace Bio-labs) and sealed with parafilm to prevent evaporation.

*LLPS and starburst reversibility experiments using temperature.* A647-labeled A1PrD (1:300 labeled to unlabeled ratio; 67 nM A647-labeled:20 μM unlabeled A1PrD) with 10 μM RNA in 200 mM NaCl, αβγ buffer was pipetted onto a MatTek glass bottom dish (MatTeK Corporation) pre-coated with Sigmacote (Sigma-Aldrich). Paraffin oil was added to prevent evaporation of the sample. The dish was incubated for ~2 min at temperatures from 25° to 60°C with 5°C-interval under thermal control of Tokai Hit ThermoPlate (ThermoFisher Scientific). The heating rate was about 6-12 sec/°C. The sample was then cooled back to 25°C. Different aged samples were prepared (1 hr and 4 hr, RT) and subjected to the same heating and cooling process described above to check reversibility of the condensed droplets or the aged starbursts.

For the timelapse video of A1PrD temperature reversibility (movie S11), A488-labeled A1PrD (1:300 labeled to unlabeled ratio; ~70 nM A488-labeled:20 μM unlabeled A1PrD) with 10 μM RNA in 200 mM NaCl, αβγ buffer was applied to a CELLview glass bottom slide (Greiner bio-one) pre-coated with Sigmacote (Sigma). Paraffin oil was added to prevent the evaporation of sample. The dish was incubated at 25°C for 3 min under thermal control of Tokai Hit ThermoPlate. The sample was heated up to 60°C, which took 90 sec. After an additional 210 sec-incubation, sample was cooled down to 25°C in 90 sec. The temperature for each recorded image was noted.

*LLPS and starburst reversibility experiments using 1,6-hexanediol.* A647-labeled A1PrD (1:300 labeled to unlabeled ratio; 67 nM A647-labeled:20 μM unlabeled A1PrD) was combined with 3 μM ThT and 10 μM RNA in 200 mM NaCl, αβγ buffer in a Grid-50 glass bottom μ-Dish (Ibidi) pre-coated with Sigmacote. Four aged samples (1, 3, 6 and 23 hr) were prepared at RT. After incubation, hexanediol was added to a final concentration of 10% (v/v), followed by another 1 hr-incubation. Samples were imaged before and after hexanediol treatment.

*Gel and starburst reversibility experiments using 1% (w/v) SDS.* A488-labeled A1PrD (~1:300 labeled to unlabeled ratio; ~70 nM A647-labeled:20 μM unlabeled A1PrD) was prepared in αβγ buffer with various [NaCl] and [RNA] in a 384-well glass bottom plate (Greiner sensoplate plus). The samples were aged for 48 hr. Timelapse videos were immediately taken 1 min after addition of SDS (to 1% (w/v) final concentration) with 30 sec intervals for 5 min. After 90 min, the SDS solution was replaced with buffer solution containing Amytracker 480 (1:200 dilution of company stock solution; Ebba Biotech). All images collected using the spinning disk confocal microscope (Nikon).

**LLPS phase diagrams.** LLPS was quantified by UV light scattering measurements at 350 nm using Nanodrop 2000 (Thermo Fisher Scientific). Lyophilized A1PrD (20 and 40 µM final concentrations) dissolved in water were mixed with buffer containing various salt and RNA concentrations to generate the final solution conditions ([NaCl]: 0, 200, and 400 mM; and [RNA]: 0, 1, and 10 µM). UV measurements were recorded after 2-min sample incubation.

**Partition efficiency experiments.** Partition efficiency of RNA and A1PrD inside and outside condensates in selected [NaCl]-[RNA] (mM-μM) conditions (*i.e.,* 0-0, 0-10, 400-0, and 400-10) were determined from protein and RNA band intensities quantified from SDS-PAGE and Native- PAGE (without SDS in running buffer) gels, respectively. Samples were incubated for 30 min and centrifuged for 30 min. Concentrations inside condensates were calculated from band intensities of the pellet fractions. Concentrations outside condensates (supernatant fraction) were calculated from subtraction of pellet with total sample concentrations (included as reference standards). Partition efficiency was calculated as the ratio of inside vs outside condensate concentrations.

**Quantification of soluble and insoluble (hexanediol- and SDS-treated) fractions.** 20 µM A1PrD in various solution conditions ([NaCl]: 0, 200, and 400 mM; and [RNA]: 0, 1, and 10 µM) were prepared and incubated for 30 min and 48 hr at RT. *Quantification of soluble fractions:* after initial 30 min incubation, the supernatant was obtained after 30 min centrifugation at 17,000xg and mixed with equal volume of SDS loading dye (2% SDS). After heating at 95°C for 5 min, samples were loaded onto 4-20% pre-cast Mini-PROTEAN Tris-Glycine gel (TG; Bio-Rad) and electrophoresed for 25-45 min at 160 mV RT in 1x TGS buffer (vWR). *Quantification of hexanediol- and SDS-insoluble fractions:* samples after 48 hr incubation were mixed with equal volume of 20% hexanediol (in water) or 2% SDS (with 20% glycerol, 20 mM Tris, pH 8). The mixtures were incubated for 15 min at RT and centrifuged for 30 min at 17,000xg. The pellets were dissolved in 2X SDS loading buffer (2% SDS), heated at 95°C for 5 min, and vortexed. Samples were run in SDS-PAGE gels as described above. Band intensities were quantified using ImageLab software (Bio-Rad).

**Image analysis.** Analysis and processing of imaging data were performed using ImageJ (NIH), Zen (Zeiss), Imaris (Bitplane) and NIS elements (Nikon) softwares. Droplets imaged at early timepoints (within 30 min) were analyzed for size and aspect ratios using the Analyze Particles Plugin in ImageJ. Other measurements such as filament lengths and thickness, droplet core diameters, intensity sum and surface volumes were analyzed using Imaris. Movies of raw data and surface rendered images were generated using Imaris. Normalized ThT intensity (sum of fluorescence intensity divided by the surface volume) of the aging droplets were fitted to an exponential model {y = y0 + A*e^(-x/t^_1/2_^)^}, where y0 is the normalized ThT intensity at time 0 and A is the signal amplitude.

**Fluorescence Recovery After Photobleaching (FRAP).** FRAP imaging was performed using a Zeiss LSM880 laser-scanning confocal microscope system with a 40x objective for selected sample conditions using 200 nM A647-labeled:20 μM unlabeled A1PrD and A488-labeled f1 RNA in different [NaCl]-[RNA] (mM-μM) conditions (*i.e.,* 0-10, 400-0, and 400-10). Different regions of interest (ROI; ~2 μm diameter spots) were selected, and the reference ROIs were drawn in adjacent regions. Following 2-3 baseline images, ROIs were bleached for 200 iterations at 100% laser power (633 and 488 nm) and were imaged for up to 4-6 min post-bleaching to calculate fluorescence recovery. FRAP recovery curves were corrected for background photobleaching (reference ROIs) and normalized against pre-bleach intensity values.

**Transmission electron microscopy (TEM).** Negative-stain TEM images were collected using a Hitachi H7500 microscope with 80-kV accelerating voltage, equipped with an AMT XR-16 digital camera at 4,000-40,000X magnification. The 300-mesh copper grids (FCF300-Cu, Electron Microscopy Sciences) were incubated with protein or protein-RNA samples in various conditions (with 10% w/v PEG-8K or 10 µM f1 RNA) at varying timepoints (30 min to 19 hr). For the temperature-induced aggregation experiments, the mesh grids were soaked with the protein samples in a PCR tube and incubated at 55°C using a PCR machine (ProFlex, ThermoScientific) for 1 and 4 hr, respectively. Afterwards, the grids were washed three times by soaking in distilled water for 30 sec, and then negatively stained with 1% uranyl acetate for 0.5 -3 min, followed by three times washing with distilled water. Additional TEM images were taken using JEOL 1400+ TEM and imaged at 80 kV using an AMT XR16 mid-mount camera. Aged samples (48 hr) with different NaCl and RNA concentrations were dropped onto formvar and carbon coated copper slot grids and allowed to sit for 2 min.  The grids were wicked off with lint- free filter paper.  Sample grids were negatively stained using 2% aqueous phototungstic acid by the drop method onto each grid and allowed to sit for 2 min.  The grids were wicked off again with lint-free filter paper.  Grids were allowed to dry overnight.

**FLIM instrumentation and experiments.** Fluorescence Lifetime Imaging Microscopy (FLIM) experiments were conducted at pH 7.5 in RT (~21.5 ± 1°C) using a custom-built ISS Alba confocal laser microscopy system (ISS) that employed an Olympus IX81 microscope equipped with an Olympus 60X/1.2 NA water objective lens, galvo-controlled mirrors, and imaging and FastFLIM^TM^ modules (ISS). Instrument calibration was carried out using rhodamine 110 dye in water (4.0 ns lifetime). Samples were prepared as described above for general fluorescence imaging measurements. For the different NaCl and RNA combination experiments, 1:200 labeled to unlabeled ratio (200 nM A488-labeled:20 μM unlabeled A1PrD) was used. Frequency domain FLIM data was analyzed using phasor analysis(*5*) and by FD lifetime fitting (fig. S3). For FD fitting, FLIM images were fitted with binning size of 3x3 pixels to 5 different modulation frequencies. An example of a FLIM FD fit and corresponding residual plot is shown in fig. S3. The laser power utilized is typically within 5-50 µW. Measured fluorescence lifetimes were found to be independent of the laser powers used (5-50 µW, fig. S4). We also checked that the lifetime values are independent on binning size and z-slices (fig. S5). Confocal FLIM images and phasor plots were generated using the PhasorAnalysis module in the VistaVision software V4.2 (ISS). FLIM histograms were generated using VistaVision V4.2 (ISS) and sometimes re-plotted using the OriginPro 9.8.5.212 software.

*Generating phasor plots:* Phasor analysis is a means of presenting fluorescence decay as pixels of an image. Using phasor representation, different molecular species can be observed through the clustering of pixels on specific regions of the phasor plot. Briefly, data at each pixel is transformed through equation E1:

$g_{i,j}\left( \omega\right)=m\cos(\varphi)$

$s_{i,j}\left( \omega\right)=m\cos(\varphi)$

where at each modulation frequency (ω), FLIM data consists of both phase delay (φ) and amplitude modulation ratio (m) where the indexes i and j identify a pixel of the original image.

The values of this transformation are then plotted on a two-dimensional histogram, or phasor plot, where each pixel of the FLIM image is represented as a point in the plot. For single-lifetime species, lifetime can be directly transformed by equation E2:

$g_{i,j}=1/(1+\omega^{2}\tau_{i,j}^{2})$

$s_{i,j}=1/(1+\omega^{2}\tau_{i,j}^{2})$

where τ is the frequency decay of the species. The two coordinates of a phasor representing a single-lifetime species has a relationship:

$${(g-0,5)}^{2}+ s^{2}=0.25$$

From the phasor plot, the lifetime of a species can be determined by its coordinate value.

*Quantification of fluorescence lifetime clusters:* FLIM images were fitted with binning size of 3x3 pixels to 5 different modulation frequencies to obtain fluorescence lifetimes for the various conditions. For the dilute conditions (non-LLPS conditions), images were taken with the absence of unlabeled protein. For different [NaCl]-[RNA] (mM-μM) conditions (*i.e.,* 0-0, 0-10, 400-0, and 400-10), fluorescence lifetimes were calculated from the region averaged values considering only the condensed phases (intensities >300 au). For RNA- and PEG-modulated LLPS conditions, fluorescence lifetimes were calculated from multiple images grouped and classified based on their incubation time and morphology. For PEG-induced LLPS: early droplets (10-30 min incubation), aged droplets (>30 min), and starbursts (>5-24 hr, and presence of filaments). For RNA-enhanced LLPS: early droplets (10-30 min incubation), initiating or early starbursts (<3 hr and presence of filaments), and late starbursts (>5-24 hr and presence of filaments). To generate statistics from multiple heterogeneous images, fluorescence lifetimes for each group were clustered into four lifetime bins: <2.1, 2.1-2.45, 2.45-2.8 and >2.8 ns. The bins were based on observed distinct peaks for representative morphologies (Fig. 3 and 5, fig. S9-11 and 16-18).

**Disorder prediction and pI calculation.** hnRNPA1 amino acid sequences correlating to NTD, RRM1, RRM2, and CTD were submitted to ProtParam (*6*) where theoretical isoelectric points were computed. Domain disorder scores were determined from IUPRED(*6-8*).

**Supplementary References**

1.	D. G. Gibson *et al.*, Enzymatic assembly of DNA molecules up to several hundred kilobases. *Nat Methods* **6**, 343-345 (2009).

2.	A. C. Ferreon, Y. Gambin, E. A. Lemke, A. A. Deniz, Interplay of alpha-synuclein binding and conformational switching probed by single-molecule fluorescence. *Proc Natl Acad Sci U S A* **106**, 5645-5650 (2009).

3.	A. C. Ferreon, J. C. Ferreon, P. E. Wright, A. A. Deniz, Modulation of allostery by protein intrinsic disorder. *Nature* **498**, 390-394 (2013).

4.	A. Molliex *et al.*, Phase separation by low complexity domains promotes stress granule assembly and drives pathological fibrillization. *Cell* **163**, 123-133 (2015).

5.	M. A. Digman, V. R. Caiolfa, M. Zamai, E. Gratton, The phasor approach to fluorescence lifetime imaging analysis. *Biophys J* **94**, L14-16 (2008).

6.	E. Gasteiger *et al.*, ExPASy: The proteomics server for in-depth protein knowledge and analysis. *Nucleic Acids Res* **31**, 3784-3788 (2003).

7.	Z. Dosztanyi, Prediction of protein disorder based on IUPred. *Protein Sci* **27**, 331-340 (2018).

8.	G. Erdos, Z. Dosztanyi, Analyzing Protein Disorder with IUPred2A. *Curr Protoc Bioinformatics* **70**, e99 (2020).
